# Novel SLAMF1-derived peptide induces apoptosis in multiple myeloma cells by targeting IRF4 transcription factor for degradation

**DOI:** 10.1101/2025.05.07.652607

**Authors:** Ingvild Bergdal Mestvedt, Eline Menu, Kristine Misund, Tobias S. Slørdahl, Kashif Rasheed, Camilla Izabel Wolowczyk, Hanne Hella, Liv Ryan, Therese Standal, Terje Espevik, Maria Yurchenko

**Author notes:** **Corresponding author**: Yurchenko Maria, **Address**: NTNU, Department of Clinical and Molecular Medicine (IKOM), P.O. Box 8905, 7491 Trondheim, Norway, **E-mail:**.

## Abstract

Multiple myeloma (MM) is the second most common hematological cancer. It remains incurable, highlighting the urgent need for novel therapeutic targets and treatment strategies. In this study, we investigated a panel of peptides derived from a functional motif of Signaling Lymphocytic Activation Molecule Family 1 (SLAMF1) with single amino acid substitutions and found that several of them exhibited potent anti-cancer activity by inducing MM cell death. The most potent peptide P7N4 significantly reduced the viability of both IL-6-dependent and -independent human myeloma cell lines (HMCLs), proteasome inhibitor-resistant MM cells, and primary MM cells, while exerting minimal effects on healthy blood cells. Furthermore, P7N4 enhanced the efficacy of the chemotherapeutic agent melphalan. When combined with the proteasome inhibitor bortezomib, P7N4 potentiated the anti-cancer effect of bortezomib in a highly aggressive murine MM model. Mechanistically, P7N4 induced apoptosis in MM cells by disrupting key pro-survival pathways, leading to reduction in IRF4, MYC, and β-catenin levels, as well as inhibition of Akt and ERK1/2 phosphorylation. Furthermore, in peptide-sensitive HMCL, P7N4 significantly altered the expression of IRF4-associated genes. These effects were likely mediated by direct interaction of peptide with IRF4, targeting this transcription factor for degradation. Overall, our findings established P7N4 as a promising therapeutic candidate for MM, warranting further optimization and in-depth mechanistic studies.

## Introduction

Multiple myeloma (MM) ranks as the second most common hematological cancer after non-Hodgkins lymphoma.^1^ MM pathogenesis is highly complex, and is encompassed by changes in gene expression, significant genomic instability, alterations in the bone marrow microenvironment, changes in cytokine and chemokine expression, and deregulation of multiple signaling pathways.^2^ The transcription factor cellular myelocytomatosis oncogene (*MYC*) is one of the major drivers of MM pathogenesis.^3^ Specific *MYC* activation signature is shown to be overactivated in 65% of MM cases, but not in normal plasma cells.^4^ Another MM-associated transcription factor is interferon regulatory factor 4 (IRF4). IRF4 is overexpressed in MM patients compared to healthy PCs and serves as a prognostic marker, with lower expression linked to longer survival.^5^ Dysregulation of the above mentioned signaling pathways and transcription factors leads to increased MM proliferation and survival, and resistance to therapy.^2^

In the recent decade, treatment of MM has substantially evolved, and includes several lines of treatment: alkylating agents, proteasome inhibitors, immunomodulatory drugs, inhibitors of nuclear export monoclonal antibodies and, most recently, bi-specific antibodies and CAR-T therapies.^6,7^ However, the high prevalence of drug resistance leads to relapse in most patients, ultimately resulting in disease progression and death.^1^ Thus, there is an urgent need to develop novel approaches and identify new therapeutic targets in MM.

Anti-cancer peptides, usually <50 amino acids in length, represent a novel and promising cancer treatment approach.^8^ Several peptides have already demonstrated the potential to inhibit cancer cell proliferation and metastatic capacity.^8^ The advantages of peptides include little or no organ accumulation, high specificity and selectivity, high biocompatibility, cost-effectiveness, and ease of modification.^9^ With the discovery of cell-penetrating peptides as a tool for intracellular delivery, it became possible to design peptides that could target specific protein-protein interactions (PPIs) inside the cancer cells.^9,10^

Signaling lymphocytic activation molecule family 1 (SLAMF1) is an immunoglobulin-like receptor with important co-stimulatory functions in hematopoietic cells. ^11^ SLAMF1 is a member of a SLAM family of type I transmembrane receptors expressed by normal immune cells, and by malignant cells of lymphoid origin, like MM, cutaneous T-cell lymphomas, B-cell non-Hodgkins lymphoma, chronic lymphocytic leukemia and Hodgkins lymphoma.^11–13^ We have recently shown that SLAMF1 amplifies signaling downstream of Toll-like receptor 4 (TLR4) by interaction with Toll-like receptor adaptor molecule (TRAM) and blocking TRAM recruitment to TLR4.^14^ We developed a synthetic peptide, P7, derived from the signaling domain of SLAMF1 and linked to the cell-penetrating peptide Penetratin (Pen) to facilitate intracellular delivery. Our findings show that P7 strongly inhibits the production of pro-inflammatory cytokines (e.g., IL-6 and TNF) induced by the TLR4-NF-κB, prevents endotoxin shock-mediated death and exhibits cardio protection after myocardial infarction in murine disease models.^15,16^

TLR4 and downstream nuclear factor kappa B (NF-κB) are known to be involved in promotion of tumor cell growth, survival, proliferation, invasion, migration and stem cell maintenance.^17,18^ Thus, targeting TLR4-NFκB cascade may be highly relevant for MM treatment, and we hypothesized that the P7 peptide could inhibit proliferation and survival of MM cells. Here, we demonstrate that while treatment of MM cancer cells with the original P7 peptide did not alter their viability, the modified peptide P7N4, with a single amino acid substitution (Y4N), exhibited a significant inhibitory effect on MM cell survival and enhanced the cytotoxic effects of several conventional MM drugs both *in vitro* and *in vivo*. We hypothesize that these properties of the P7N4 peptide are TLR4- independent and linked to the direct inhibition of IRF4, leading to the suppression of pro-survival signaling pathways and apoptosis in MM cells.

## Materials and methods

### Primary cells

The study was performed in accordance with the Helsinki declaration. Informed consent was obtained from all patients or donors, the use of primary cells was approved by the Regional Committee for Medical and Health Research Ethics (#2009/2245, #2009/2029). Human PBMCs were isolated from buffy coats (St. Olavs Hospital, Trondheim, Norway) as previously described.^19^ B cells from PBMCs were isolated by negative selection (B Cell Isolation Kit II, Milentenyi Biotech, Cologne, Germany). PBMCs, B cells and murine 5T33MMvt cells were cultured in RPMI1640 (Merck, Schnelldorf, Germany) supplemented with 10% fetal calf serum (FCS) from Gibco (Life Technologies, Oslo, Norway), 100 U/ml penicillin and 100 μg/ml streptomycin (pen/strep) (Life Technologies). Patient samples were obtained were collected by the Norwegian Myeloma Biobank (Biobank1, St. Olavs Hospital). CD138^+^ and CD138^-^ were isolated using RoboSep automated cell separator and human CD138 positive selection kit (StemCell Technologies, Cambridge, UK; Cat: 17877RF). Cells were maintained in RPMI1640 supplemented with 2% heat-inactivated (HI) human serum, pen/strep and 1 ng/ml rhIL-6 (Gibco).

### Cell lines

HMCLs used in this study include ANBL-6 (from Dr. Diane Jelinek, Mayo Clinic, Rochester, MN, USA), JJN3 (from J. Ball, University of Birmingham, UK), INA-6 (from Dr. Gramatzki, Erlangen, Germany), RPMI-8226, U266 (ATCC, Rockville, MD, USA), OH-2, IH-1, KJON (in-house) and AMO-1 (from Dr. Christoph Driessen Le, Switzerland). The proteasome inhibitor-resistant cell lines AMO-BTZ, AMO-CFZ and INA-6 BTZ described in ^21–23^. Cell lines, if not indicated otherwise, were cultured in RPMI1640 containing 10% FCS. ANBL-6 and INA-6 were supplemented with 1 ng/ml rhIL-6, while IH-1, KJON and OH-2 were supplemented with 10% HI human serum and 2 ng/ml rhIL-6. RPMI-8226 and U266 were supplemented with 20% and 15% FCS, respectively. The primary murine MM cells Vk*MYC (12653) were a kind gift from Marta Chesi, Mayo clinic.^20^ Vk*MYC cells were kept in RPMI1640 with L-Glut, 2% HI human serum, pen/strep and 1 ng/ml rhIL-6. All media variants contained pen/strep.

### Peptides, reagents

Synthetic peptides were obtained from GenScript (Rijswijk, Netherlands) or Life Technologies. Peptide modifications and characteristics were as previously described.^15^ LPS (ultrapure LPS *E. coli* strain K12, ultrapure LPS *E. coli* strain 0111:B4) obtained from was obtained from InvivoGen (San Diego, CA, USA). Melphalan (Merck, Darmstadt, Germany) was dissolved in 90% ethanol and 2.775% hydrochloric acid. Proteasome inhibitors carfilzomib and bortezomib are from Selleck Chemicals (Houston, TX, USA). Inhibitors for caspase-8 (Z-IETD-FMK), caspase-9 (Z-LEHD-FMK) were from Biotechne (Oslo, Norway).

### Cell viability assay

The cell viability was evaluated using Cell Titer-Glo 2.0 Assay (Promega, Germany) according to the manufacturer’s instructions. Prior assay, cells were seeded in 96-well plates with 100 μl/well of media containing 5000 cells/well for HMCLs, 5T33vt, 10000 cells/well for Vk12653, 20000 cells/well for primary B cells and CD138^+^ primary MM cells, 50000 cells/well for PBMCs and CD138^-^ cells. Luminescence was recorded with POLARstar Omega (BMG LABTECH, Mölndal, Sweden) using integration time of 1 sec/well. Three to five technical triplicates were done for all treatments. Standard curve for the ANBL6 cell line was prepared by serial dilutions (1:2). The relative values for percentage of viable cells were calculated using relative luciferase units (RLU) for control or other treatments. The standard curve and values based on standard curve were calculated using microplate manager 6 (MPM6) software (BioRad, California, USA).

### Flow cytometry

Flow cytometry was performed as previously reported ^22^ using Annexin V/PI staining (Apoptest Annexin AV-FITC kit, NeXins Research, Kattendijke, Netherlands). Briefly, 50000 cells/well for AMO-1 or primary CD138^+/-^ cells in 200 μl/well of media were seeded in round-bottomed 96-well plates (NUNC) and incubated with control treatment or peptide as indicated in the result section and figure legends. The cell suspension was subsequently transferred to pre-cooled flow tubes with PBS, centrifugated (1500 rpm, 5 min, 4 °C) and stained with Annexin V (AV) and propidium iodide. Unstained and single stained samples were used as controls. Evaluation of Vk12653 cancer cells content was performed as described.^24^ Data was acquired using a LSRII flow cytometer (BD Biosciences, Franklin Lakes, NJ, USA) and analyzed in FlowJo 10.8.1 sofware (Tree Star, Ashland, OR, USA).

### Cell lysis, pull-downs by biotinylated peptides and western blotting

For signaling studies, HMCLs (2.5 mill/ml) were treated in cell culture flasks, followed by cell lysis, SDS-PAGE and western blotting. At each timepoint, cells were washed by cold PBS before adding lysis buffer: 1X RIPA lysis buffer (150 mM NaCl, 50 mM Tris-HCl, pH 7.5, 1% Triton X-100, 5 mM EDTA, 50 mM NaF, 2 mM Na_3_VO_4_, protease inhibitors (Thermo Fisher Scientific), and phosphatase inhibitors (Roche)) or 1X NP40 lysis buffer (150 mM NaCl, 50 mM Tris-HCl pH 7.5, 0.5% IGEPAL CA-630 (Merck), 1 mM EDTA, protease and phosphatase inhibitors). Cells were lysed for 30 min at 4 °C with agitation and cleared by centrifugation. Protein concentrations were measured using Pierce^TM^ BCA Protein Assay Kit (Thermo Fisher Scientific) according to the manufacturers protocol, with 10-50 ug of samples used for SDS-PAGE. Pulldown (PD) assays by biotinylated peptides were performed as previously described.^15^ Co-precipitated proteins in PDs were eluted with 1× loading buffer (Thermo Fisher Scientific) containing 40 mM DTT (Merck) and heating of samples, followed by SDS-PAGE and WB. SDS-PAGE and Western blotting are described in detail in ^15^. Lower parts of gels for PDs were stained by SimplyBlue Safe Stain (Thermo Fisher Scientific) to access peptides’ loading. Images of stained gels were taken on Gel Doc EZ Imager (Bio-Rad, Bio-Rad Laboratories), and densitometry analysis of stained bands was performed using Image Lab Software 6.0.1 (Bio-Rad).

### Antibodies

The following primary antibodies were used for western blotting: mouse anti-β-catenin (Cat: 610153) from BD Biosciences; rabbit anti-PARP-1 (#ab32138), β-tubulin (#ab6046), mouse anti-GAPDH (#ab9484) from Abcam (Cambridge, UK); rabbit anti-GSDMD (#97558), phospho-MLKL (#91689), caspase-9 (#9502), cleaved caspase-8 (#9496), cleaved caspase-3 (#9664), cleaved caspase-7 (#94915), MYC (#5605), phospho-Akt Ser473 (#4060), pERK1/2 Thr202/Tyr204 (#9101), IRF4 (#4299) and mouse anti-β-tubulin (#86298) were from Cell Signaling Technology (Danvers, MA, USA). Secondary antibodies (HRP-linked) swine anti–rabbit (#P039901-2) and goat anti-mouse (#P044701-2) were from Dako Agilent Technologies (Santa Clara, CA, USA).

### RNA sequencing

AMO-1 and RPMI-8226 cells (2 x 10^6^ cells/sample) were treated by 10 μM P7N4-Pen peptide, or media, for 90 min followed by RNA isolation and bulk RNA sequencing. Total RNA was isolated using the RNasy Mini Kit (Qiagen, Hilden, Germany). Bulk RNAseq was performed at the Genomics Core facility (NTNU, Norway) using the TruSeq Stranded mRNA Library Prep Kit (Illumina, San Diego, CA, USA) and 400 ng input RNA followed by 75 bp single read sequencing on the Illumina Hiseq 4000. FASTQ files quality controlled with fastqc (v0.11.9) then filtered and trimmed by fastp (v0.20.0). Trimmed sequences were aligned to the genome reference using STAR (v2.7.3) and quality metrics were extracted with picard CollectRNASeqMetrics (v2.21.5). Transcript counts were generated using quasi alignment (Salmon v1.3.0) to the GRCh38 transcriptome reference sequences. Transcript counts were imported into the R statistical software and aggregated to gene counts using the tximport (v1.14.0) bioconductor package for downstream statistical analysis. Gene counts were normalized and analyzed for differential expression using the DESeq2 bioconductor package. Heatmap was generated using free online tool (heatmapper.ca).

### Animal studies

All animal experiments were performed in compliance with the ARRIVE guidelines, procedures approved by the Ethical Committee for Animal Experiments of the Vrije Universiteit Brussel (project number: 23-281-16). C57BL/KalwRij female mice (Envigo Laboratories, Horst, Holland), age 4-6 weeks, were housed in animal facility 3 mice/cage. Mice were acclimatized and aged for 9 weeks prior to the start of the experiment. Mice were injected i.v. on day 0 with 5 x 10^5^ 5T33MMvv cells in 200 μl PBS and divided into four groups (n = 12): vehicle (sterile water), single agent groups BTZ or P7N4-Pen and combination of P7N4-Pen with BTZ. Starting from day 1, mice were treated i.p. five times per week either with 10 mg/kg P7N4-Pen or vehicle for the designated treatment groups. Starting from day 2, 0.6 mg/kg bortezomib was given twice per week i.p. Injection volumes did not exceed 100 µl. At day 21, mice were euthanized by cervical dislocation following bleeding by retro-orbital puncture. Post-mortem, mice spleens were harvested and weighed, the percent plasma cells in the BM was determined on May Grunwald-Giemsa-stained BM smears from one femur, and the M-component was measured as previously described.^25^

### Statistical analysis

Number of experiments and statistical test are indicated in figure legends. Results were analyzed using the GraphPad Prism 10.1.2 (GraphPad Software Inc, La Jolla, CA, USA).

## Results

### Modified SLAMF1-derived peptides reduce the viability of MM cells

The effect of SLAMF1-derived peptides on MM cell viability was assessed using the CellTiter-Glo assay. ANBL-6 cells, one of the most representative MM models,^26^ were treated with a panel of peptides for 48 h, which included the previously described P7 peptide^15^ and peptides with single amino acid substitutions in the P7 sequence (Fig. 1A, 1B; Supplementary Fig. 1A, 1B, see Additional file 1). The screen identified five peptides with significant inhibitory effect on cell viability: P7K2-Pen, P7A4-Pen, P7N4- Pen, P7D4-Pen and P7S4-Pen (Fig. 1A, 1B). All five peptides exhibited a dose-dependent decrease in MM cell viability, whereas the original P7-Pen peptide and the control peptide Pen showed no such effect (Fig. 1C). P7N4-Pen demonstrated the strongest inhibitory effect on cell viability, both at 24 (Supplementary Fig. 1C) and 48 h of treatment (Fig. 1C). Using a standard curve generated by serial dilution of ANBL-6 cells, we confirmed that P7N4-Pen reduced the number of viable cells per well to levels significantly below the initial seeding number, indicating that the peptide induces cell death (Supplementary Fig. 1D). A time-course evaluation revealed that P7N4-Pen inhibitory impact on cell viability was comparable at 24 and 48 h and significantly increased at 72 h (Fig. 1D).

**Fig. 1:**
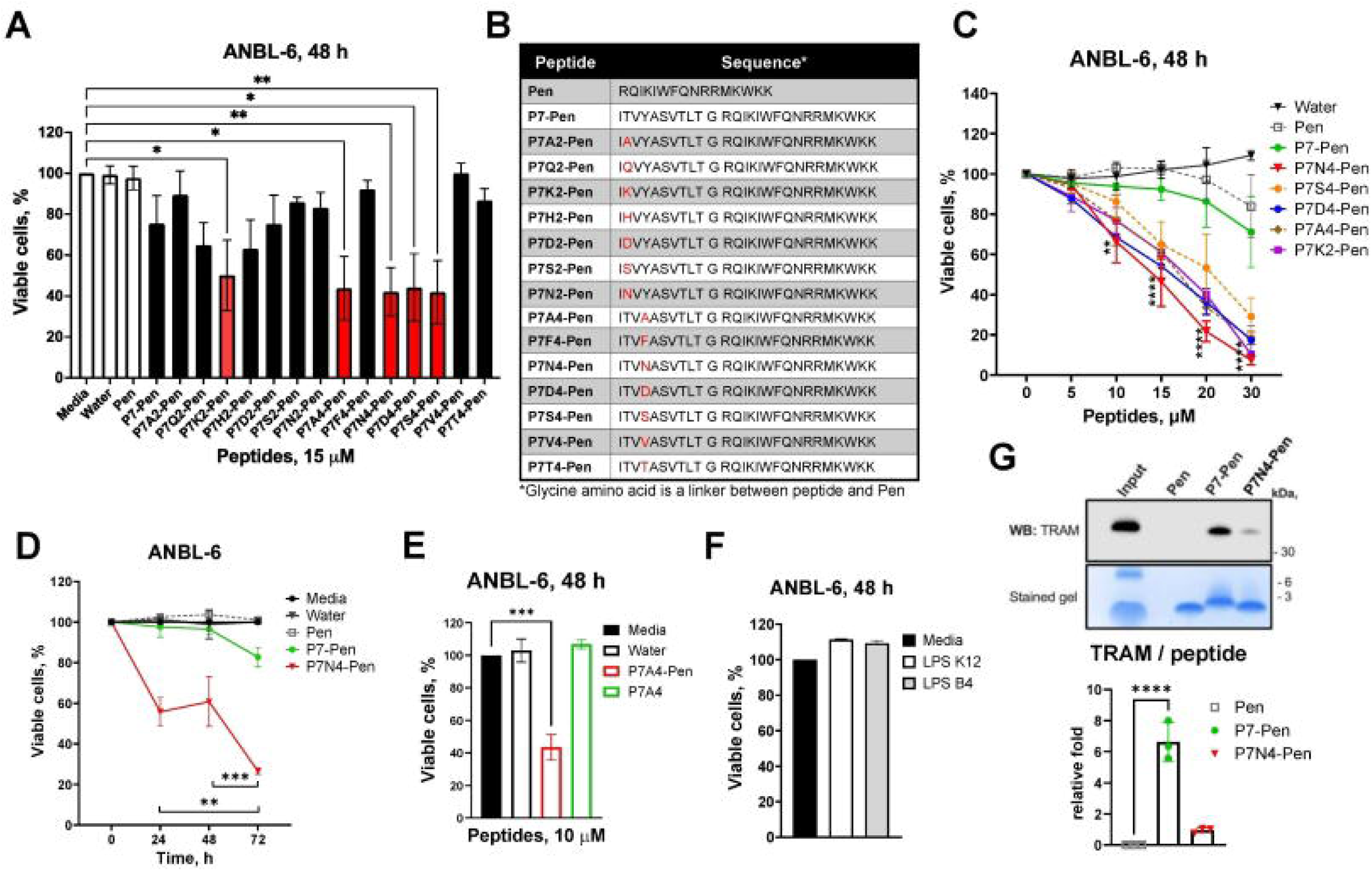
SLAMF1-derived peptides significantly reduce the viability of ANBL-6 cells independently of TLR4 signaling inhibition. (**A**) Cell viability assay for ANBL-6 cells treated for 48 h with media containing water (solvent) or 15 μM peptides - control peptide penetratin (Pen) or SLAMF1-derived peptides – P7-Pen and its variants with single amino acid substitutions at positions 2 and 4. (**B**) Table with sequences for peptides used in the screen, with amino acid substitutions in relation to P7-Pen peptide highlighted in red. (**C**) ANBL6 cells were treated with increasing concentrations of peptides or water control for 48 h, followed by cell viability assay. Significance is shown for P7N4-Pen *vs.* water. (**D**) ANBL-6 cells were treated with 10 μM P7N4-Pen for 24, 48 or 72 h before accessing cell viability. Significance is shown for P7N4-Pen effect between different time points. (**E**) ANBL-6 cells were treated for 48 h with P7A4-Pen or P7A4, or control treatments, followed by cell viability assays. (**F**) Results of cell viability assay for ANBL-6 cells grown media with 1 μg/ml K12 or O1111:B4 LPS for 48 h. (**E**) Western blot analysis for TRAM on pulldowns from ANBL-6 lysates by biotinylated Pen, P7-Pen or P7N4-Pen peptides immobilized on streptavidin beads. Pen-biotin was used as a negative control. Whole-cell lysate was loaded as input control (7.5% from the total sample used for pulldown). Lower part of the gels was stained by SimplyBlue SafeStain for peptides’ loading control. Ratio for TRAM signal to the intensity of peptides’ staining was quantified for three independent experiments and presented as mean ± SD. (**A, C, D-F**) Data presented as the mean± SEM of viable cells when related to the media control (100%) for three or four independent experiments. Significance evaluated using (**A, D, E**) one-way ANOVA, Dunnett’s multiple comparisons test; (**C**) two-way ANOVA, Tukey’s multiple comparisons test; significance levels: *P\**<0.05, *P\*\**<0.01, *P\*\*\**<0.001, *P\*\*\*\**<0.0001). Only significant results are shown.

To confirm that attachment to Pen for intracellular entry was the pre-requisite for the peptide’s efficacy, we compared the effect of peptides with and without Pen tag. Due to solubility issues, we were unable to use P7N4 without CPP, so we tested another peptide from the primary screen - P7A4. While P7A4-Pen significantly reduced MM cell viability, the P7A4 peptide was completely ineffective (Fig. 1E). These results confirm that the peptide should be tagged with Pen for intracellular delivery to preserve its efficacy.

Stimulation of ANBL-6 cells with 1 μg/ml of ultrapure LPS (K12 or O111:B4) for 48 h induced only a marginal increase in proliferation (Fig. 1F), suggesting low impact of TLR4-mediated signaling on proliferation in these cells. Unlike P7-Pen, P7N4-Pen did not co-precipitate TRAM adaptor protein (Fig. 1G), which is a direct target of P7-Pen in the TLR4 signaling pathway.^15^ Altogether, our findings suggest that the inhibitory effect of P7N4-Pen on MM cell viability is TLR4-independent.

### P7N4-Pen reduces viability of both IL-6 dependent/independent HMCLs

Most HMCLs are derived from cells outside the bone marrow and are independent of its microenvironment.^27^ To enhance their relevance as models, IL-6-dependent cell lines were developed.^28^ Since IL-6 receptor signaling may influence treatment response,^29^ we compared P7N4-Pen’s effect on IL-6-dependent (Fig. 2A) and IL-6-independent HMCLs (Fig. 2B-D). P7N4-Pen peptide significantly reduced the viability of seven out of the eight tested HMCLs – INA-6, IH-1, OH-2, KJON, JJN3, U266 and AMO-1 (Fig.2A-C; Supplementary figure 2A, 2B, see Additional file 2). The only resistant cell line was the IL-6 independent RPMI-8226 (Fig. 2D). P7N4-Pen dose-dependently decreased the viability of AMO-1 and INA-6 cell lines, with IC_50_ being in the range of 4-8 μM (Fig. 2C; Supplementary Fig. 2A). Overall, P7N4-Pen exhibited a significant inhibitory effect on viability of most of the tested HMCLs, independent of IL-6 requirement.

**Fig. 2:**
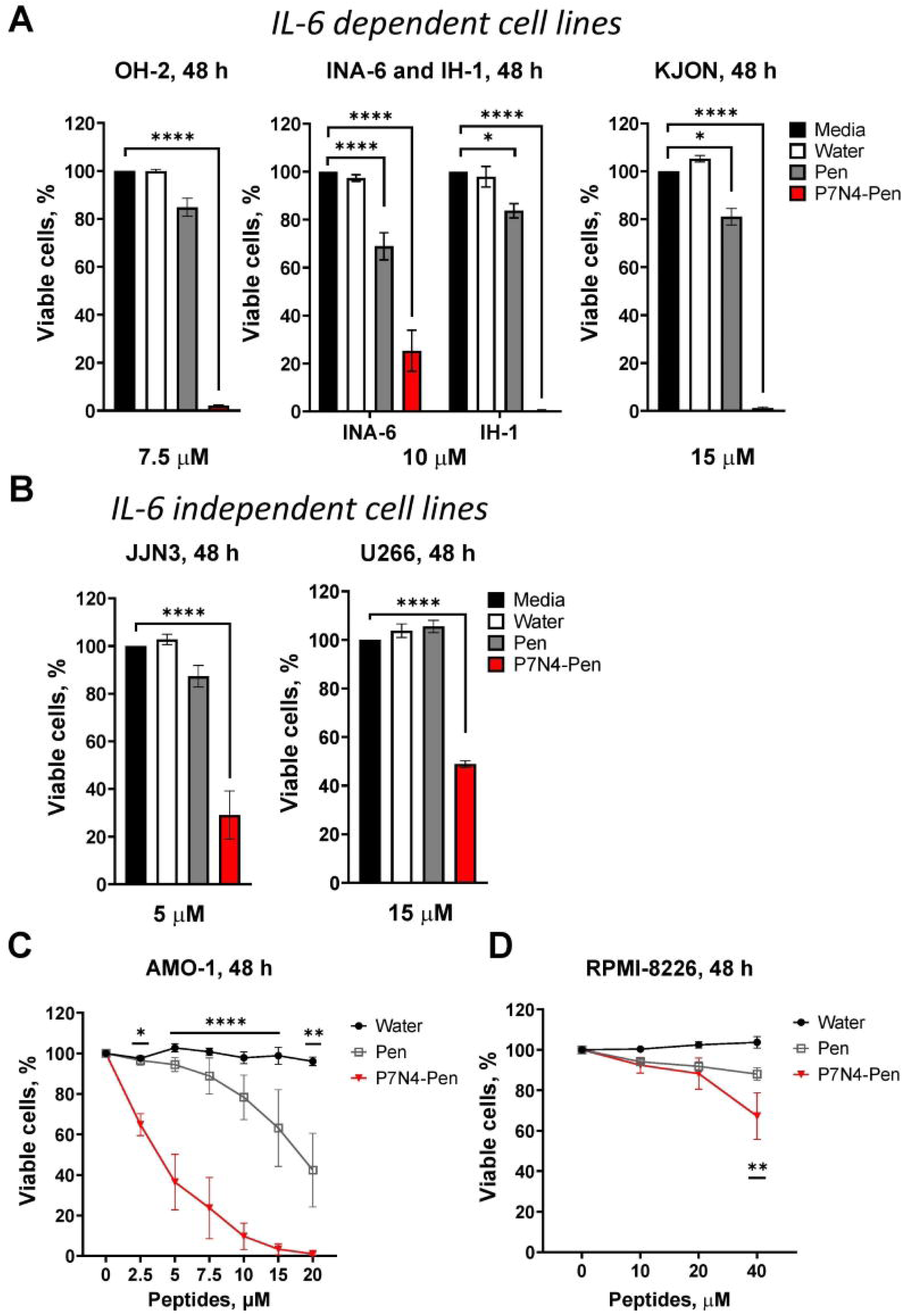
P7N4-Pen significantly reduces the viability of both IL-6-dependent and IL- 6-independent HMCLs. (**A**) IL-6 dependent (INA-6, IH-1, OH-2, KJON) and (**B, C, D**) IL-6 independent (JJN3, U266, AMO-1, RPMI-8226) HMCLs have got P7N4-Pen (concentrations indicated in the graphs) or control treatments (media, water or control peptide Pen) and were incubated for 48 h before measuring cell viability by CellTiterGlo assay. Data presented as the mean number of viable cells when related to the media control (100%) with mean ± SEM for three biological replicates. (**C, D**) Significance was evaluated between Pen or P7N4-Pen treated samples. Significance evaluated using two-way ANOVA, Dunnett’s multiple comparisons test, significance levels: *P\**<0.05, *P\*\**<0.01, *P\*\*\**<0.001). Only statistically significant results are shown.

### P7N4-Pen demonstrates selective cytotoxicity toward CD138^+^ MM primary cells and specificity for cancer cells over normal cells

Further we evaluated the peptide’s effect on the viability of primary MM cells. We have used freshly isolated CD138^+^ from bone marrow aspirates of MM patients, which were treated with 15 μM peptides for 18 h (Fig. 3A; Supplementary Fig. 3A, 3B, see Additional file 3). P7N4-Pen, but not the control peptide, significantly reduced the viability of primary CD138^+^ cells in a dose-dependent manner, though the response varied among patients (Fig. 3A; Supplementary Fig. 3A).

**Fig. 3:**
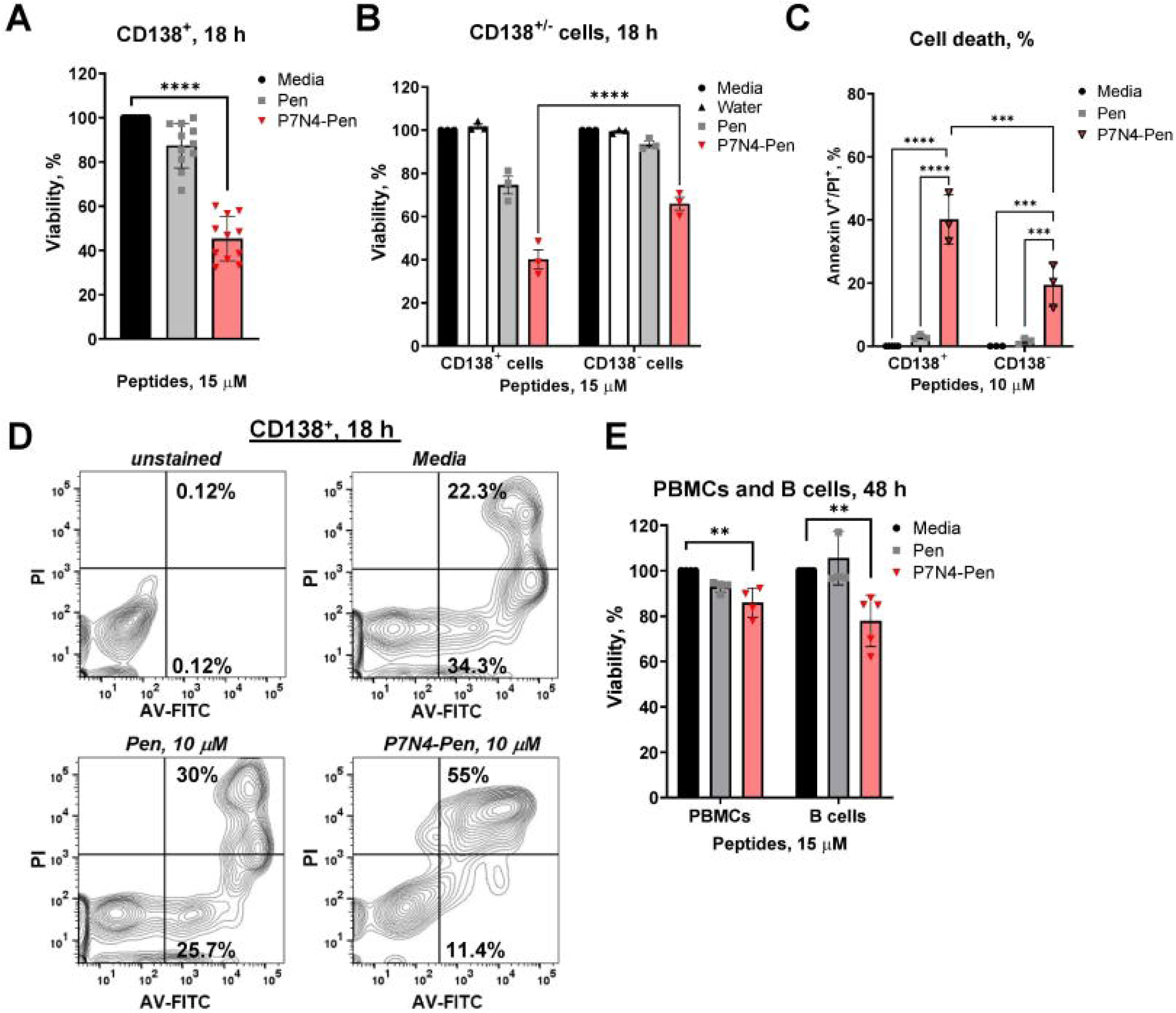
P7N4-Pen induces cell death in cancer cells with much less cytotoxic effect towards normal cells. (**A, B**) Graphs show viability of primary CD138^+^ MM cells (n = 11) and (**B**) direct comparison of viability between pre-treated primary CD138^+^ and CD138^-^ cells (n = 3) isolated from bone marrow aspirates of MM patients. Cells were incubated for 18 h with control treatment (media, water or 15 μM Pen) or 15 μM P7N4- Pen before measuring cell viability. (**C, D**) Cell death induction in CD138^+^ and CD138^-^ cells was evaluated in 18 h of treatment by peptide or control treatments by flow cytometry-based double staining assay using Annexin V-FITC (AV-FITC) and propidium iodide (PI) (n = 3). (**C**) Combined data for mean percentage of dead cells (AV^+^/PI^+^, %). (**D**) Representative flow cytometry graphs and gating. (**E**) PBMCs and B cells from healthy donors (n = 4 or n = 6, respectively) were incubated for 48 h in growth media, or media containing water or 15 μM peptides (Pen, P7N4-Pen) before cell viability was measured. (**A-B, E)** Cell viability assessed with CellTiterGlo assay, data presented as mean ± SEM percentage of viable cells when related to the media control (100%). Significance evaluated using (**A, E**) one-way ANOVA, Dunnett’s multiple comparisons test; (**B, C**) two-way ANOVA, Sidaks multiple comparisons test; significance levels: *P\*\**<0.01, *P\*\*\**<0.001. Only statistically significant results are shown.

To assess the peptide’s specificity towards cancer cells, we compared its effect on CD138^+^ and CD138^-^ bone marrow cells (Fig. 3B). While P7N4-Pen reduced both populations’ viability, its impact was significantly stronger on CD138^+^ cells (Fig. 3B, Supplementary Fig. 3B). The CD138^-^ cell fraction may include both normal and cancerous cells due to MM-associated CD138 loss,^30^ which could explain the peptide’s effect on CD138^-^ cells.

Reduced ATP levels in CellTiter-Glo assay can indicate both decreased metabolic activity and cell death. To confirm that P7N4-Pen induced cell death, we conducted Annexin V (AV, an early apoptosis marker) and propidium iodide (PI, a dead cell marker) staining, followed by flow cytometry (Fig. 3C, D). P7N4-Pen significantly increased the AV^+^/PI^+^ double-positive population, indicating enhanced cell death in primary MM cells compared to control treatments, with stronger effect towards CD138^+^ than CD138^-^ cells (Figure 3C).

We further tested peptide on peripheral blood mononuclear cells (PBMCs) and B cells from healthy donors (Fig. 3E). P7N4-Pen caused only a slight reduction in the viability of PBMCs and B cells (Fig. 3E), which was significantly less pronounced when compared to HMCLs (Fig. 1, 2) or primary MM cells (Fig. 3A) under similar conditions. These results suggest that P7N4-Pen selectively targets cancerous cells over healthy cells.

### P7N4-Pen enhances melphalan efficacy and induces cell death in proteasome inhibitor-resistant HMCLs

Alkylating agents are often associated with severe side effects, as seen with high-dose melphalan,^31^ making dose reduction a priority to improve patient quality of life. Here, we investigated whether P7N4-Pen could enhance melphalan’s therapeutic effects, potentially enabling a reduction in its dosage without compromising efficacy. Using two P7N4-sensitive cell lines - JJN3 (IL-6 independent) and ANBL-6 (IL-6 dependent) - we found that P7N4-Pen significantly augmented melphalan’s inhibitory effects. The combined treatment with the peptide and melphalan was significantly more effective than melphalan alone at all tested concentrations (Fig. 4A, B). Furthermore, P7N4-Pen enhanced melphalan-induced cytotoxicity in CD138^+^ primary cells (Fig. 4C; Supplementary Fig. 4A, see Additional file 4). These findings suggest that combining P7N4-Pen with melphalan could allow for lower therapeutic doses of the alkylating agent.

**Fig. 4:**
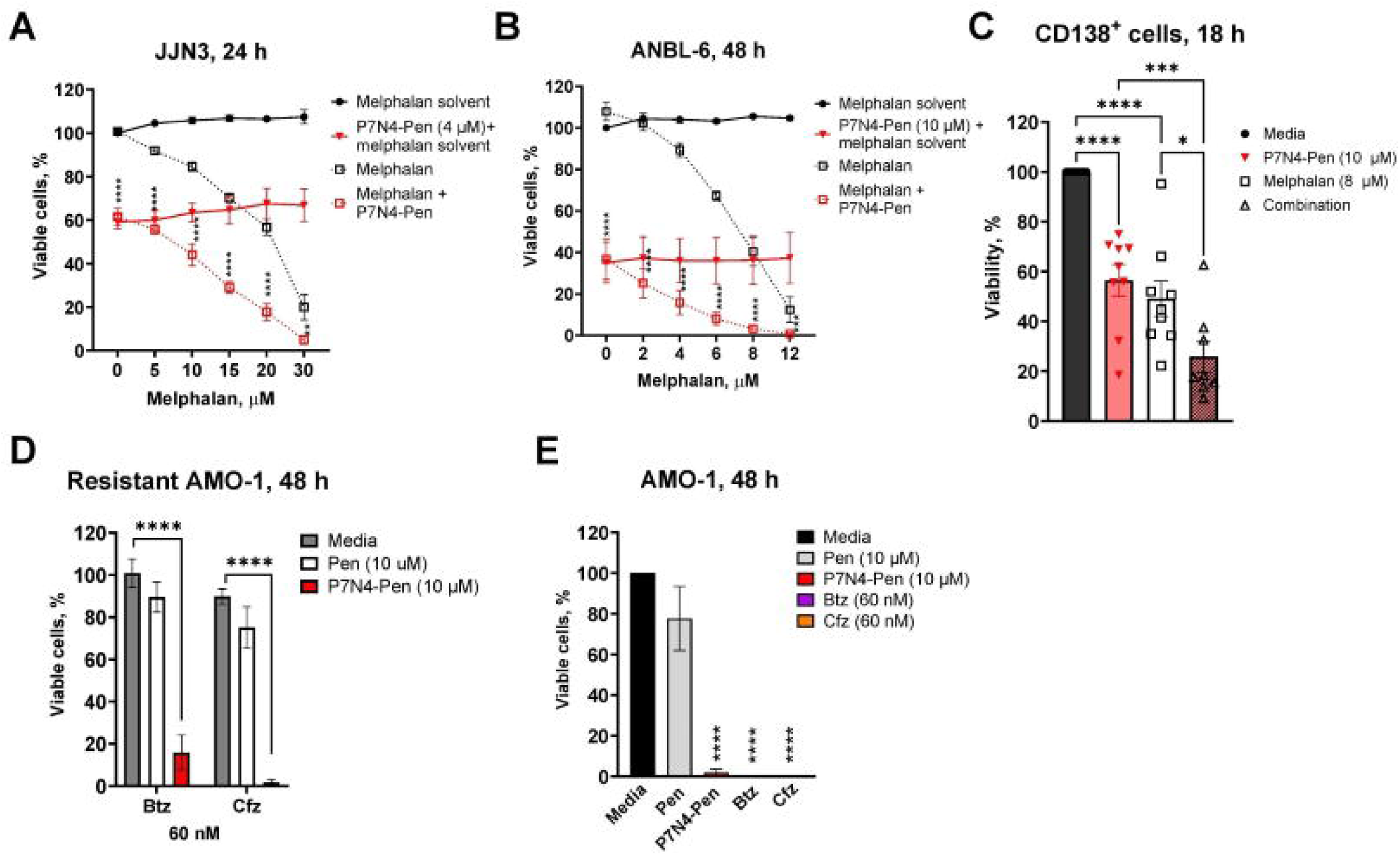
P7N4-Pen potentiates the inhibitory effect of melphalan and reduces the viability of proteasome inhibitor-resistant cells. (**A, B**) Cell viability assay for JJN3 and ANBL-6 HMCLs treated for the indicated time with melphalan solvent (EtOH), individual drugs, or combinations of solvent and P7N4-Pen, as well as melphalan and P7N4-Pen, at the concentrations specified in the graphs. Significance is shown for melphalan *vs.* combination group (melphalan and P7N4-Pen). (**C**) Combined graph showing the viability of primary CD138⁺ MM cells isolated from bone marrow aspirates of MM patients (n = 8-9). Cells were treated for 18 h with 8 μM melphalan, 10 μM P7N4- Pen, or their combination before cell viability was assessed. Cells cultured in media served as the control. (**D**) AMO-1 cells resistant to bortezomib (Btz) or carfilzomib (Ctz) were maintained in media containing 60 nM proteasome inhibitors (PIs) and treated for 48 h with 10 μM P7N4-Pen or Pen before assessing cell viability. (**E**) Viability of wild type AMO-1 cells was addressed after 48 h treatment by peptides (10 μM) or Btz and Ctz (60 nM), performed in parallel with treatment of resistant cells shown on (**D**). Cell viability was measured using the CellTiterGlo assay (ATP release). Data are presented as the percentage of viable cells relative to the solvent control (**A, B**) or media control (**C-E**) (set to 100%). Values represent the mean ± SEM from three independent experiments (**A, B, D, E**) or individual patients (**C**). Significance evaluated using (**A, B**) two-way ANOVA, Sidak’s multiple comparisons test; (**C**) One-way ANOVA, Tukeýs multiple comparisons test, (**D, E**) two-way ANOVA, Tukeýs multiple comparisons test. Significance levels: \**P*<0.05, \*\**P*<0.01, \*\*\**P*<0.001, \*\*\*\**P*<0.0001. Only statistically significant results are shown.

Next, we investigated whether resistance to proteasome inhibitors impacts sensitivity to P7N4-Pen. AMO-1 wild type and cells resistant to both bortezomib (Btz) and carfilzomib (Cfz) were incubated in media containing 60 nM Btz or Cfz, either alone or in combination with Pen or P7N4-Pen peptide (Fig. 4D, E). Proteasome inhibitors completely eliminated wild type AMO-1 cells (Fig. 4E). At the same time, P7N4-Pen significantly reduced the viability of both wild type and resistant cells (Fig. 4D, E). Similar results were obtained for Btz-resistant INA-6 cells (Supplementary Fig. 4B). Thus, P7N4-Pen has the potential to be an effective treatment if resistance to proteasome inhibitors develops.

### P7N4-Pen induces both extrinsic and intrinsic apoptosis in MM cells

We hypothesized that apoptosis is the primary mode of cell death induced by P7N4- Pen. Here, AMO-1 cells were pre-treated with peptide and stained by AV and PI, followed by flow cytometry analysis. P7N4-Pen initiated cell death as early as 4 h after treatment, inducing up to 20% of AV^+^ cells (early apoptotic) and 50% of AV^+^/PI^+^ (dead) cells (Fig. 5A, B), indicating that peptide initiates apoptosis in these cells.

**Fig. 5:**
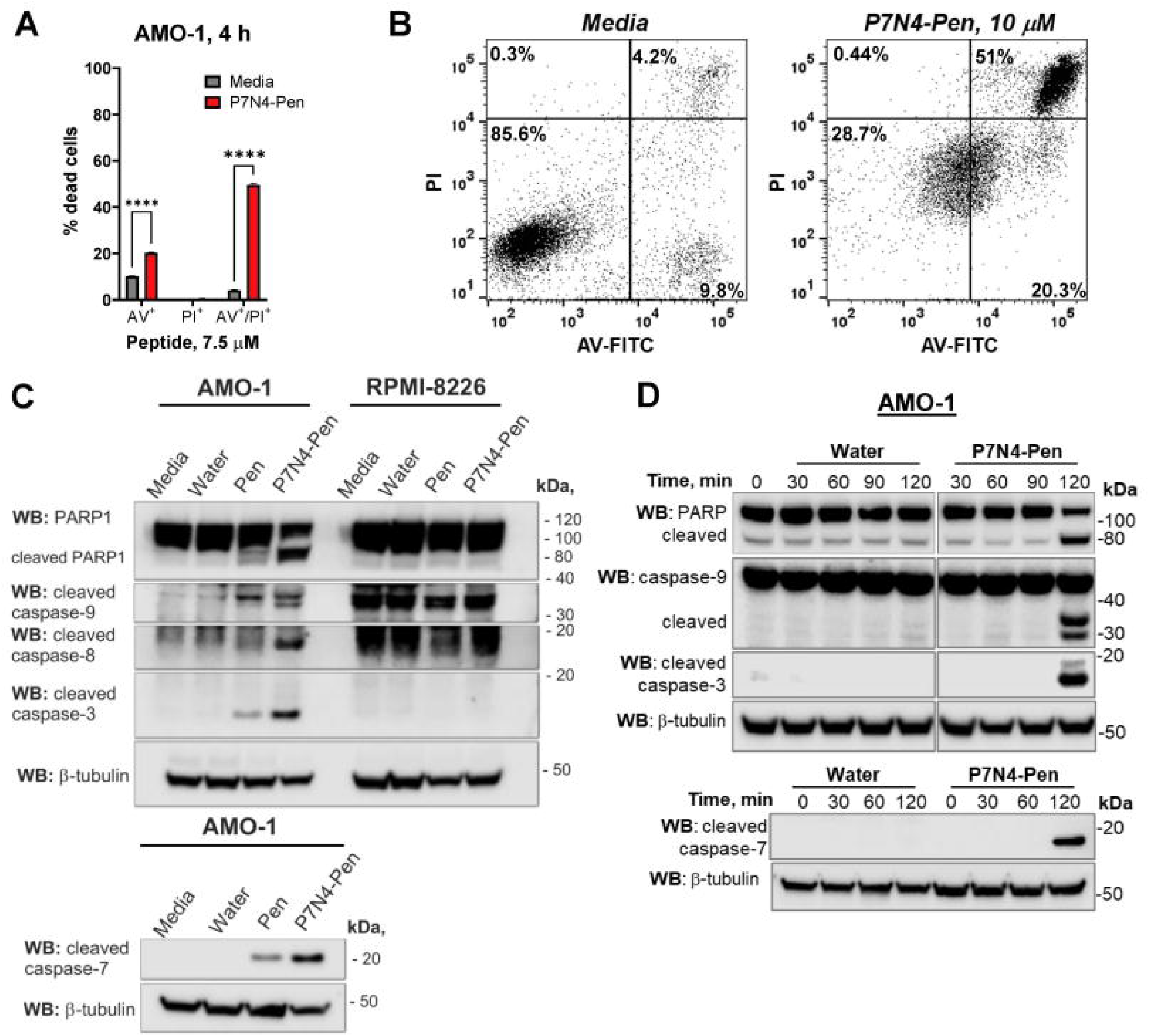
P7N4-Pen initiates apoptosis in MM cells. (**A, B**) AMO-1 cells were treated with P7N4-Pen (7.5 μM) or kept in cell culture media for 4 h prior to double staining by Annexin V-FITC (AV-FITC) and propidium iodide (PI), and flow cytometry analysis. Results from three independent experiments. Data presented as mean ± SD, statistical significance evaluated by two-way ANOVA, Sidaks multiple comparisons test, significance levels ****P<0.0001. Only statistically significant results are shown. Representative plot with gating is shown on (**B**). (**C**) AMO-1 or RPMI-8826 cells were treated for 6 h by peptides (7.5 μM for sensitive AMO-1 cells, 10 μM for resistant RPMI- 8226 cells) or kept in growth media with/without peptide solvent (water) prior WB analysis of PARP1 or caspases cleavage. (**D**) AMO-1 lysates were treated by 7.5 μM P7N4-Pen or control treatments for designated time, followed by WB analysis of PARP1 or caspases cleavage. Images for the Western blots for the selected conditions/treatments were excised from the images of the same membranes, with full image shown in supplementary materials. (**C, D**) Images for representative experiments are shown (n = 3), WB for total β-tubulin applied for loading control.

Apoptosis occurs via two pathways: caspase-8 (extrinsic) and caspase-9 (intrinsic), both converging on effector caspases (-3, -6, -7).^32^ To identify the predominant pathway induced by peptide, we analyzed cleaved caspases and caspase target protein PARP1 by Western blot. P7N4-Pen induced cleavage of caspases -8, -9, -3, -7, and PARP1 in peptide-sensitive AMO-1 cells but not in resistant RPMI-8226 cells (Fig. 5C). Caspase-3 and PARP1 cleavage were also confirmed in ANBL-6 cells (Supplementary Figs. 5A, 5B, see Additional file 5). P7N4-Pen, but not the P7-Pen peptide, induced the cleavage of caspases and PARP1 within 2 h (Fig. 5D, Supplementary Fig. 5C), consistent with no effect of P7-Pen on MM cell viability (Fig. 1A). The control peptide Pen induced minimal caspase and PARP1 cleavage along with much stronger pro-apoptotic effect of P7N4- Pen (Supplementary Fig. 5C).

P7N4-Pen did not alter MLKL Ser358 phosphorylation (necroptosis marker) or generate the 31-kDa GSDMD cleavage product (pyroptosis marker) (Supplementary Figs. 5D, 5E). Instead, it produced a 43-kDa inactive GSDMD fragment.^33^ Thus, it is likely that necroptosis or pyroptosis are not contributing to the peptide-mediated cell death.

To establish the relative contribution of extrinsic *vs.* intrinsic apoptosis pathways, we used caspase -8 and -9 inhibitors (Z-IETD-FMK and Z-LEHD-FMK, respectively). Inhibition of either caspase reduced the peptide-mediated cleavage of caspase-3 and PARP1 (Supplementary Fig. 5F), indicating that both pathways contribute to P7N4-Pen-induced apoptosis.

### P7N4-Pen inhibits key survival pathways in MM cells

Inhibiting key survival pathways, such as Wnt/β-catenin, MYC-IRF4 transcriptional interplay, and ERK1/2 and Akt activation, is a potential strategy for targeting MM cells.^2–5^

Here we show that P7N4-Pen significantly reduced β-catenin and MYC protein levels in peptide-sensitive AMO-1 and ANBL-6 HMCLs within 6 h, but not in resistant RPMI-8226 cells (Fig. 6A; Supplementary Fig. 6A).

**Fig. 6:**
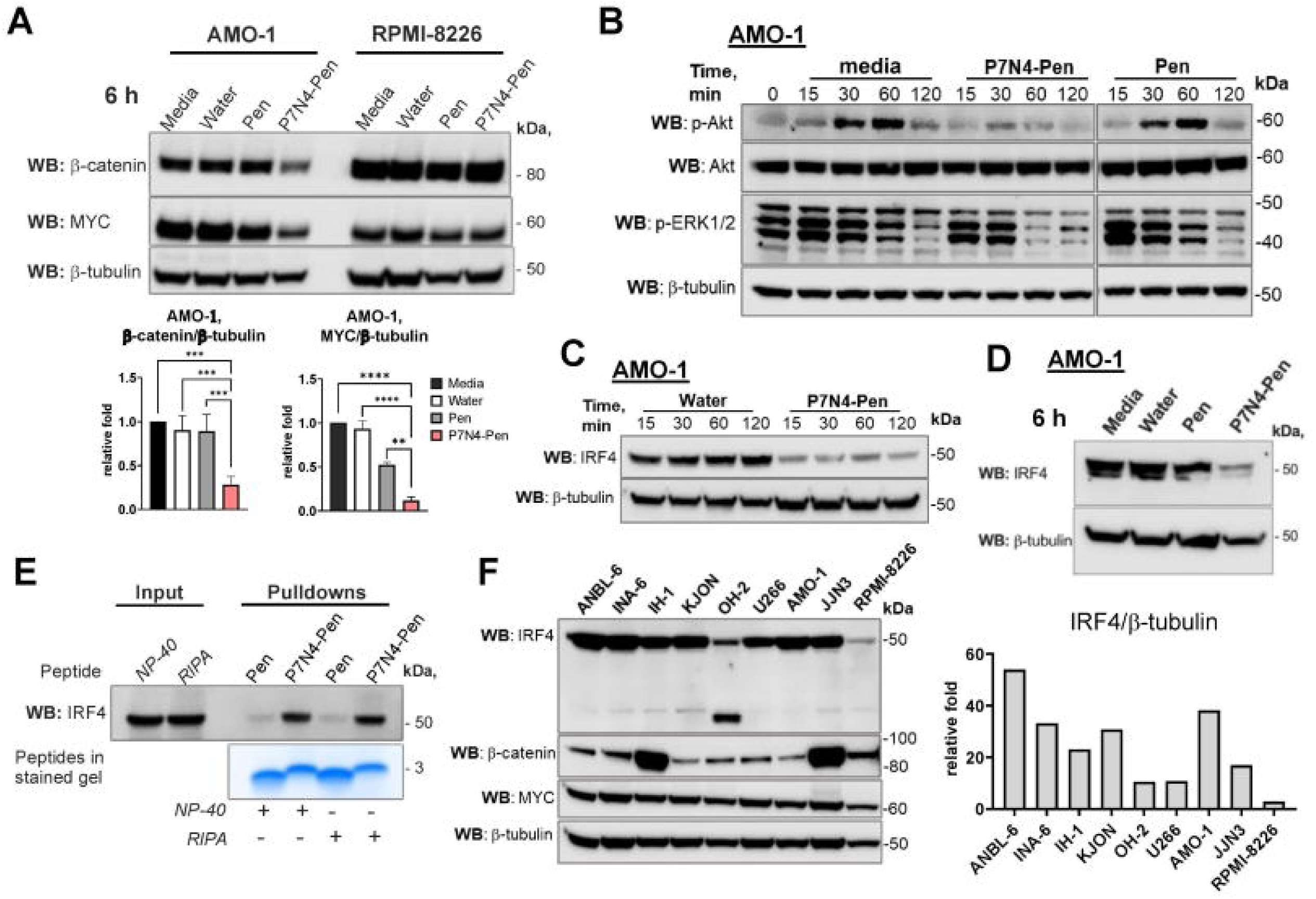
Peptide decreases IRF4, MYC and β-catenin expression, and Akt and ERK1/2 phosphorylation in MM cells. (**A**) WB analysis of total β-catenin and MYC levels in lysates of AMO-1 and RPMI-8226 cells that were treated with peptides (7.5 μM for AMO-1, 10 μM for RPMI-8226) or control treatments (media, water) for 6 h. The level of protein expression normalized to β-tubulin is shown on graph as relative fold, mean ± SD from three independent experiments, statistical analysis performed using one-way ANOVA, significance levels: \*\**P*<0.01, \*\*\**P*<0.001, \*\*\*\**P*<0.0001. (**B, C**) AMO-1 cells were resuspended in fresh culture media and immediately treated by 7.5 μM peptides for indicated time prior to cell lysis and WB analysis of phospho-Akt (S473), total Akt, phospho-ERK1/2 (Thr202/Tyr204) (**B**) and IRF4 (**C**). Images for Western blots shown on (**B**) are from the same membranes. (**D**) AMO-1 cells were treated with 7.5 μM peptides or control treatments for 6 h prior to WB analysis of total IRF4 expression. A representative image from three independent experiments is shown. (**E**) WB analysis of proteins that co-precipitated with biotinylated peptides in pulldowns from the lysates of AMO-1 cells (500 μg of protein/pulldown). Lysis buffers, NP40 or RIPA as described in methods, with modification of RIPA buffer to high salt (300 mM NaCl). Cell lysates were loaded for input control (7.5% from the sample used for pulldown). Lower part of the gel was stained by SimplyBlue SafeStain for peptides’ loading control. Representative images of one of three independent experiments are shown. (**F**) Western blot analysis of total IRF4 expression by HMCLs. Graph shows the relative fold of IRF4 expression to β-tubulin for the representative image. (**A-D, F**) WB for total β-tubulin was applied for loading control.

AMO-1 cells exhibited high basal ERK1/2 phosphorylation, and media exchange induced robust Akt phosphorylation (Fig. 6B). P7N4-Pen, but not Pen, downregulated the phosphorylation of both kinases (Fig. 6B). As Akt inhibits pro-apoptotic proteins,^34^ and MEK1/2 (upstream of ERK1/2) inhibition induces MM cell apoptosis,^35^ P7N4-Pen’s suppression of Akt and ERK1/2 phosphorylation may contribute to apoptosis.

IRF4, a key MM survival factor,^5,36^ was rapidly downregulated by P7N4-Pen within 15 min, with effects sustained at 6 h in AMO-1 and ANBL-6 cells (Fig. 6C-D, Supplementary Fig. 6C). Given this rapid decrease in IRF4 protein levels, we hypothesized that IRF4 might be a direct target of P7N4. Pulldown assays with biotinylated peptides confirmed that P7N4-Pen, but not Pen, co-precipitated with IRF4 in AMO-1 lysates under mild and stringent conditions (Fig. 6E). This suggests a direct peptide-IRF4 interaction. Interesting that RPMI-8226 cells had the lowest IRF4 expression among tested HMCLs, correlating with their weak response to P7N4-Pen, while β-catenin and MYC levels were comparable to other HMCLs (Fig. 6F).

To further explore these findings, we performed RNA sequencing (RNA seq) analysis of control or P7N4-Pen-treated AMO-1 and RPMI-8226 cells, focusing on differential expression of IRF4-associated genes,^37^ as well as *KLF2* and *MYC.*^38,39^ There was a clear effect of peptide on the expression of these genes in AMO-1 but not in resistant RPMI-8226 cells (Fig. 7A; Supplementary Tables 1, 2). *MYC* was significantly downregulated by peptide in AMO-1 cells, while *IRF4* expression remained unchanged in both cell lines (Fig. 7B; Supplementary Tables 1-3), suggesting P7N4- Pen targets IRF4 protein rather than gene expression

**Fig. 7.**
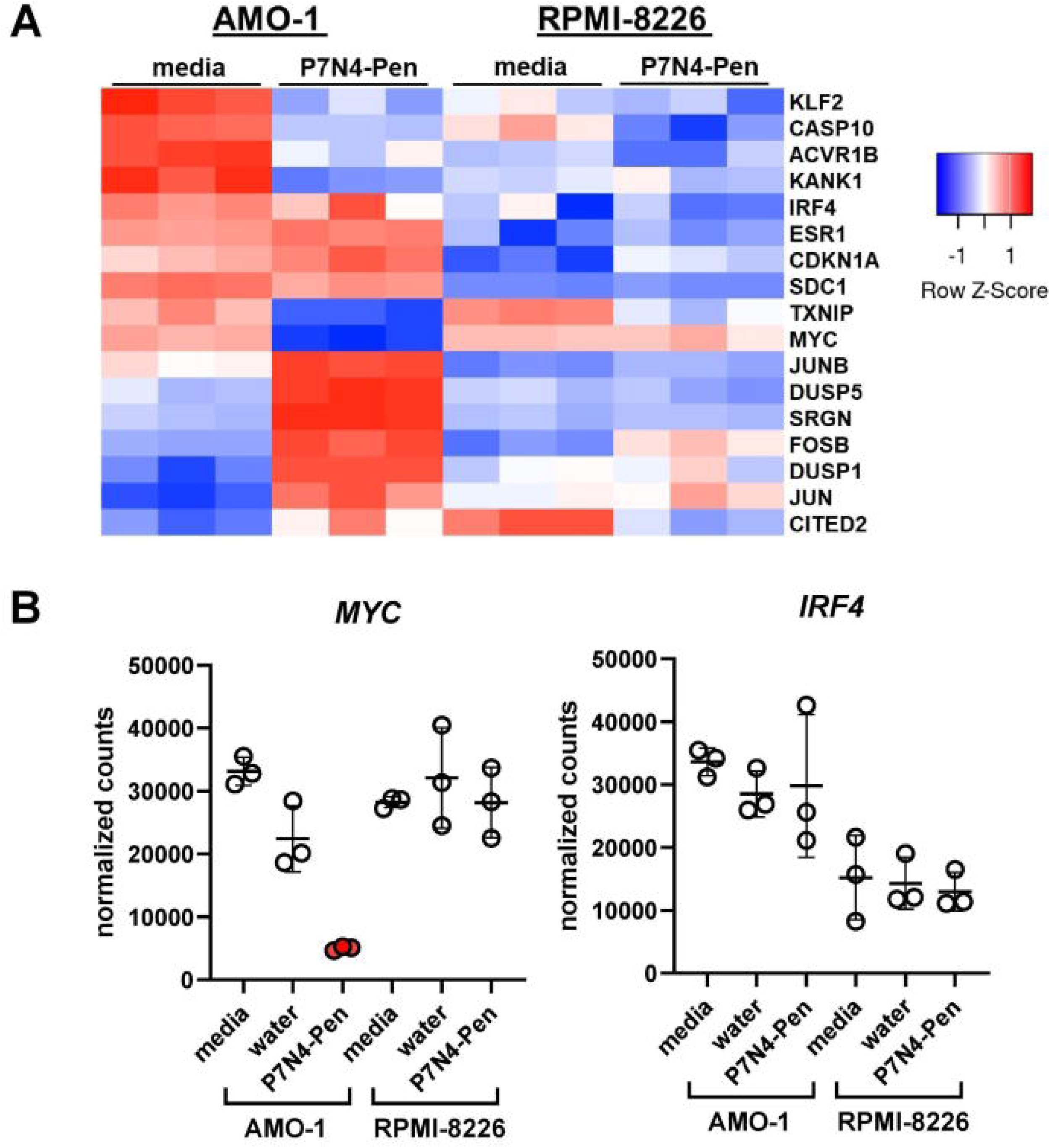
P7N4-Pen regulates the expression of IRF4-associated genes in peptide sensitive AMO-1 HMCL. AMO-1 and RPMI-8226 cells were treated with 10 μM P7N4- Pen or control treatment (media) for 90 min prior to RNA isolation and sequencing, with 3 biological replicates for each treatment condition. (**A**) Heatmap is showing gene expression levels (z-transformed log_2_ TPM) of IRF4-associated signature genes, and *IRF4*, *MYC* and *KLF2*. Genes for IRF4-associated signature were selected based on the differential analysis of their expression in AMO-1 cells by pairwise analysis for P7N4- Pen *vs.* media samples, log_2_fold change > 0.4, or log_2_fold change < -0.4, Padj < 0.05). (**B**) Graphs show normalized gene counts for *IRF4* and *MYC* in all treatment conditions for three biological replicates for each condition, mean ± SD. Normalized gene counts represent raw count matrix normalized by size factor. For statistical analysis we used raw gene counts as input, with only *MYC* mRNA expression in P7N4-Pen AMO-1 cells (red dots on *MYC* graph) being statistically significant when compared to control treatments with log_2_fold change = -2.03131976, Padj = 9.7E-15 peptide to water control, log_2_fold change = -2.636032321, Padj = 7.88E-24 peptide to media control.

### P7N4-Pen enhances anti-cancer effect of bortezomib in a highly aggressive MM in vivo model

Further we assessed the efficacy of P7N4-Pen as anti-cancer agent in two murine MM models: 5T33MMvv in C57BL/KalwRij mice (Fig. 8) and Vk*MYC (subline Vk12653) in C57BL/6 mice (Supplementary Fig. 7D, 7E). Both models mimic human MM, including proliferation in the bone marrow, serum M-component production, and bone disease.^40,41^

**Fig. 8.**
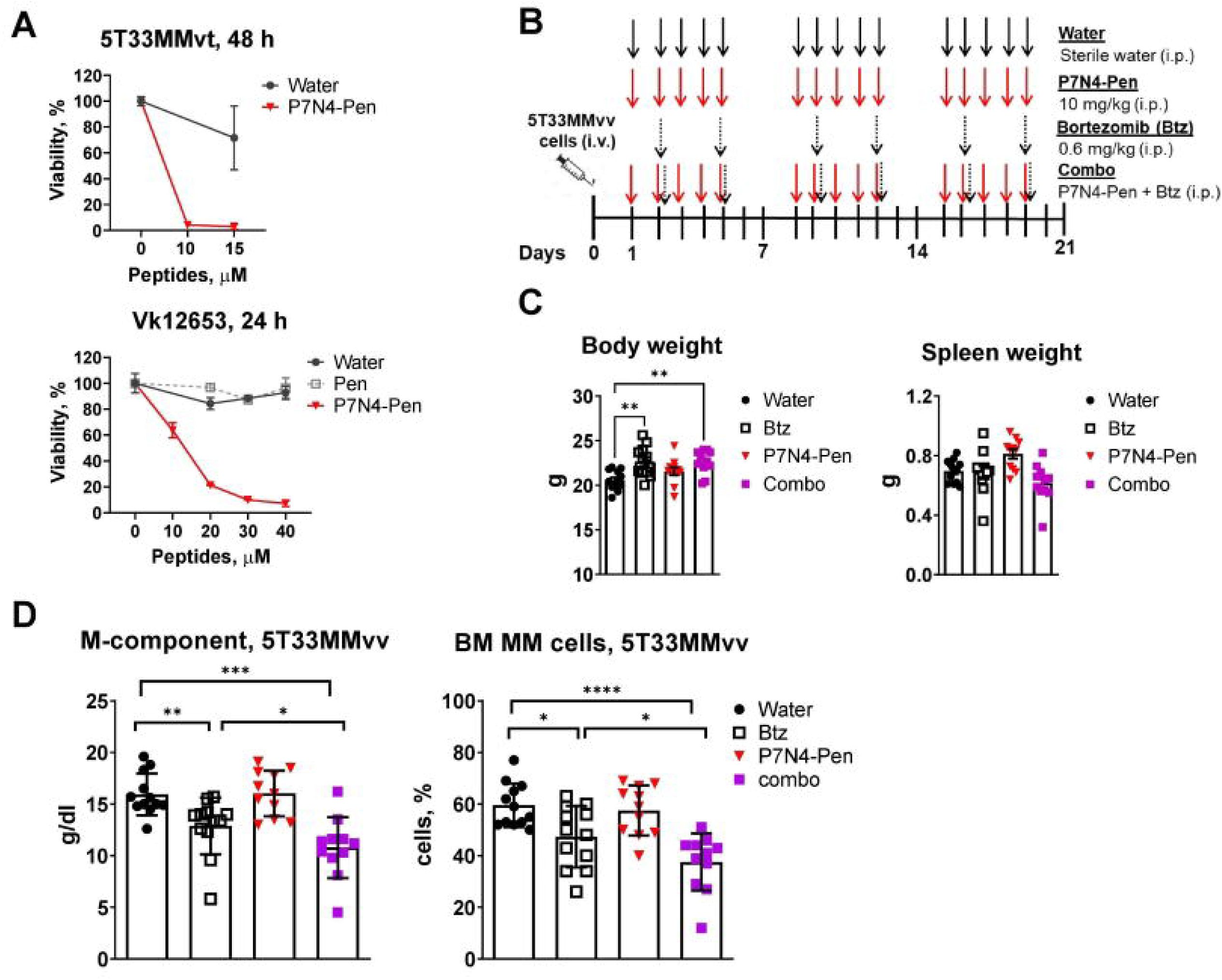
P7N4-Pen peptide potentiates the effect of bortezomib in the 5T33MMvv murine model. (**A**) Graphs depict the dose-dependent response of 5T33MMvt and Vk12653 cells to peptides, assessed by the CellTiter-Glo assay. Data are presented as the percentage of viable cells relative to the media control (set to 100%). Values represent the mean ± SD from three biological replicates. (**B**) Graphical overview of 5T33MMvv murine model setup and timing. C57BL/KaIwRij mice (n = 48) were inoculated on day 0 with 5T33MMvv cells by i.v. injection, followed by i.p. injection of peptides solvent water, or P7N4-Pen (10 mg/kg), or Btz (0.6 mg/kg) or combination of P7N4-Pen and bortezomib. Water or peptide were given 5 times a week for 3 weeks, and Btz was given twice a week for 3 weeks. Mice were euthanized on day 21, and blood and bone marrow cells were collected for analysis. (**C**) Body weight and spleen weight, and (**D**) M-component levels (g/dL) in blood samples, along with the percentage of MM tumor cells in bone marrow (quantified from May-Grünwald-Giemsa-stained BM smears on cytospins), were measured at the endpoint. (**C, D**) Data shown as mean ± SEM (n = 11-12). Significance evaluated using one-way ANOVA, Tukey’s multiple comparisons test; and for BM cells count - two-tailed Mann-Whitney test. Significance levels: *P<0.05, **P<0.01, ***P<0.001, ****P<0.0001. Only statistically significant results are shown; i.v.- intravenous, i.p.- intraperitoneal.

Evaluation of P7N4-Pen’s potential toxicity (17 mg/kg) in healthy C57BL/6 mice via repeated intraperitoneal (i.p.) injections showed a favorable safety profile, with no adverse effects on body weight and temperature, or major organs compared to control mice (Supplementary Fig. 7A-C).

*In vitro*, P7N4-Pen reduced the viability of 5T33MMvt (a stroma-independent variant of 5T33MMvv) and Vk12653 cells, with greater sensitivity in 5T33MMvt (Fig. 8A). *In vivo*, 5T33MMvv mice received water (solvent), P7N4-Pen (10 mg/kg), bortezomib (Btz), or combination (combo) treatment following the treatment timeline shown in Figure 8B. At day 21 (end point), serum and tissue samples were analyzed. While body weight was higher in Btz and combo groups, spleen weight showed no major differences (Fig. 8C). Serum M-component levels and MM cell burden in bone marrow significantly decreased in Btz and combo groups, with the combo treatment being the most effective (Fig. 8D). Overall, P7N4-Pen alone showed no significant effect but enhanced bortezomib’s efficacy.

In the Vk12653 model, cancer cells were injected intravenously (i.v.) into C57BL/6 mice five weeks before treatment initiation (Supplementary Fig. 7D). Given lower P7N4- Pen sensitivity in Vk12653 cells *in vitro* (Fig. 8A), a 17 mg/kg dose was used. Five weeks post-inoculation, mice with M-component levels (globulin-to-serum albumin ratio, G/A) between 0.35 and 0.5 were assigned to treatment groups (water control and P7N4-Pen). At the study endpoint, no significant impact on tumor burden was observed based on serum M-component, spleen weight, or MM cell percentage (Supplementary Fig. 7E).

## Discussion

To date, no peptides targeting intracellular protein-protein interactions have been clinically approved for cancer, but their therapeutic potential is immense.^42^ Features such as high specificity, strong binding affinity, targeting of “undruggable” proteins, ease of modification, high biocompatibility, and low toxicity contribute to the significant promise of peptides.^43^ For example, the PCNA-targeting peptide ATX-101 showed a favorable safety profile in cancer patients.^44^ Minimal or no side effects have also been reported for other peptides, such as Fexapotide (Phase III trials for prostate cancer).^45^ SLAMF1-derived peptides also showed a favorable safety profile in murine models, both as a single injection,^15,16^ and after repeated doses (evaluated in this study).

This study examined the cytotoxic potential of SLAMF1-derived peptides against MM cells. The original P7 peptide, a 10-amino acid fragment from the SLAMF1 receptor, inhibits TLR4-mediated inflammation *in vitro* and *in vivo.*^15,16^ As TLR4-NF-κB signaling is a potential therapeutic MM target, its inhibition could reduce the inflammatory environment supporting myeloma growth and apoptosis resistance.^17,46^ However, TLR4 inhibitor P7-Pen did not affect MM cell viability. Instead, SLAMF1-derived peptides with single amino acid substitutions (positions 2 or 4) significantly reduced MM cell viability, with P7N4-Pen being the most potent. P7N4-Pen did not co-precipitate with main P7- Pen target - TRAM adaptor protein, suggesting its anti-cancer effect is TLR4- independent, likely arising from interaction with alternative targets. Selected for further evaluation, P7N4-Pen exhibited specific cytotoxicity against most tested HMCLs (IL-6- dependent and independent) and primary MM cells, while minimally affecting healthy cells.

Advances in MM treatments have improved the five-year survival rate,^47^ yet treatment-related side effects and acquired resistance remain challenges.^48^ Bortezomib was the first proteasome inhibitor approved for the treatment of MM,^49^ representing a major breakthrough in MM therapy. Unfortunately, the patient’s cancer cells eventually become refractory to bortezomib.^50^ Here, we found that proteasome inhibitor-resistant MM cells remained susceptible to peptide treatment. *In vivo*, while P7N4-Pen was ineffective as monotherapy, it enhanced the anti-tumor effects of bortezomib, suggesting its potential as an additive therapy to eliminate resistant tumor clones.

Since the introduction in 1958, melphalan is still commonly used for treatment of MM,^51,52^ but its side effects - including bone marrow suppression, nausea, vomiting,^31^ and increased susceptibility to infections^53^ – limits its application. Identifying alternatives or dose-reduction strategies is critical to improve patient quality of life. Here, we demonstrate that P7N4-Pen may significantly enhance melphalan’s therapeutic effect, potentially allowing a substantial dose reduction.

A common feature of bortezomib- and melphalan-resistant MM cells is the acquisition of altered DNA repair mechanisms.^54^ Targeting these pathways offers a promising therapeutic strategy for relapsed and refractory MM, and potentially for newly diagnosed MM. P7N4-Pen induces cleavage of PARP1, a DNA repair protein, which may contribute to peptide’s potency in bortezomib-resistant cells and ability to enhance melphalan’s efficacy.

We investigated key aspects of the mechanism underlying peptide-induced cell death in MM cells and found that P7N4-Pen significantly downregulates the protein levels of IRF4, MYC, and β-catenin, and inhibits activation of ERK1/2 and Akt kinases, ultimately leading to apoptosis. Furthermore, P7N4-Pen co-precipitates with IRF4. IRF4 in known to bind to the *MYC* promoter region and transactivate its expression, creating a positive autoregulatory feedback loop.^36^ IRF4 is considered an Achilles’ heel in MM, as its downregulation leads to reduced cell viability, cell cycle arrest, and apoptosis.^36^ Based on our results, we propose that the binding of P7N4-Pen to IRF4 is a crucial initial step that disrupts IRF4 protein interactions, which triggers transcriptional changes in IRF4- associated genes, including the downregulation of *MYC* at both mRNA and protein levels, and ultimately leads to peptide-induced apoptosis.

Currently, researchers are exploring small molecules that may facilitate IRF4 degradation, such as proteolysis-targeting chimeras (PROTACs) designed to bind to the PU.1 interaction site in IRF4, recruiting cereblon to induce proteolytic degradation of IRF4 (conference abstract).^55^ However, the full report on this study is not yet available. Notably, P7N4-Pen is the first IRF4-degrading molecule that has already been extensively evaluated in MM cell lines, primary MM cells, and murine models.

It is also important to discuss the *in vivo* performance of P7N4-Pen. Despite its inhibitory effect on the viability of Vk12653 and 5T33MMvt murine cells *in vitro*, no such effect was observed *in vivo*. This lack of *in vivo* efficacy may be attributed to several factors. First, peptide-based drugs often have limited efficacy due to their short half-life, particularly in small rodents. It is likely that P7N4-Pen does not reach the tumor in sufficient concentrations when administered via intraperitoneal injections, suggesting that continuous infusion may be required to enhance its efficacy. Additionally, further chemical modifications could improve the therapeutic potential of P7N4-Pen. Second, both Vk12653 and 5T33MMvv models are highly aggressive and considered the challenging MM models to treat.^20,56^ In the 5T33MMvv model, an initial reduction in tumor cell count (up to fivefold) does not necessarily affect tumor burden at the endpoint,^56^ complicating drug efficacy evaluation. Vk12653 tumors exhibit dysregulated MYC expression and activity,^40^ meaning that MYC activity in this model may not be significantly dependent on IRF4. Indeed, a much higher concentration of P7N4-Pen was required to kill VK12653 cells *in vitro* compared to 5T33MMvt cells. MYC can be targeted either directly using the small molecule 10058-F4^57^ or indirectly through proteasome inhibitors.^3^ We propose that a combination therapy simultaneously targeting IRF4 with the P7N4-Pen peptide and MYC with the aforementioned or novel compounds could be a promising therapeutic strategy for MM treatment.

Overall, our study identifies P7N4-Pen as a potent degrader of IRF4, selectively inducing tumor cell death while having minimal effect towards normal cells. We suggest that further chemical modifications could enhance the peptide’s stability, targeted delivery, and efficacy *in vivo*. Overall, our findings highlight P7N4-Pen as a promising therapeutic candidate for MM and potentially other malignancies driven by aberrant IRF4 activity.

## Acknowledgements

We thank Knut Sverre Grøn, Nils Hagen and Mari Meslo Lien from the Comparative medicine Core Facility (CoMed, NTNU) for technical assistance with animal experiments. RNAseq was performed by the Genomics Core facility (NTNU, Trondheim, Norway), and we thank Arnar Flatberg from the Core facility for RNAseq data analysis.

## Author contributions

MY: conceptualization, resources, supervision, funding acquisition, project administration. TS, TE: conceptualization, funding acquisition, supervision, methodology, editing. IBM, MY: formal analysis, methodology, investigation, writing, editing. TSS: methodology, patients’ samples, resources, editing. EM: methodology, editing. KM, EM, KR, CIW, HH, LR: investigation.

## References

1. Pinto V, Bergantim R, Caires HR, Seca H, Guimarães JE, Vasconcelos MH. Multiple Myeloma: Available Therapies and Causes of Drug Resistance. Cancers (Basel). Feb 10 2020;12(2)doi:10.3390/cancers12020407

2. Hu J, Hu W-X. Targeting signaling pathways in multiple myeloma: Pathogenesis and implication for treatments. Cancer Letters. 2018/02/01/ 2018;414:214–221. 10.1016/j.canlet.2017.11.020

3. Holien T, Våtsveen TK, Hella H, Waage A, Sundan A. Addiction to c-MYC in multiple myeloma. Blood. 2012;120(12):2450–2453. doi:10.1182/blood-2011-08-371567

4. Chng WJ, Huang GF, Chung TH, et al. Clinical and biological implications of MYC activation: a common difference between MGUS and newly diagnosed multiple myeloma. Leukemia. Jun 2011;25(6):1026–35. doi:10.1038/leu.2011.53

5. Bai H, Wu S, Wang R, Xu J, Chen L. Bone marrow IRF4 level in multiple myeloma: an indicator of peripheral blood Th17 and disease. Oncotarget. 2017;8(49)

6. Berdeja JG, Madduri D, Usmani SZ, et al. Ciltacabtagene autoleucel, a B-cell maturation antigen-directed chimeric antigen receptor T-cell therapy in patients with relapsed or refractory multiple myeloma (CARTITUDE-1): a phase 1b/2 open-label study. Lancet. Jul 24 2021;398(10297):314–324. doi:10.1016/S0140-6736(21)00933-8

7. Morè S, Corvatta L, Manieri VM, Morsia E, Poloni A, Offidani M. Novel Immunotherapies and Combinations: The Future Landscape of Multiple Myeloma Treatment. Pharmaceuticals (Basel). Nov 19 2023;16(11)doi:10.3390/ph16111628

8. Xie M, Liu D, Yang Y. Anti-cancer peptides: classification, mechanism of action, reconstruction and modification. Open Biology. 2020;10(7):200004. doi:doi:10.1098/rsob.200004

9. Araste F, Abnous K, Hashemi M, Taghdisi SM, Ramezani M, Alibolandi M. Peptide-based targeted therapeutics: Focus on cancer treatment. Journal of Controlled Release. 2018/12/28/ 2018;292:141–162. 10.1016/j.jconrel.2018.11.004

10. Cabri W, Cantelmi P, Corbisiero D, et al. Therapeutic Peptides Targeting PPI in Clinical Development: Overview, Mechanism of Action and Perspectives. Review. Frontiers in Molecular Biosciences. 2021-June-14 2021;8doi:10.3389/fmolb.2021.697586

11. Farhangnia P, Ghomi SM, Mollazadehghomi S, Nickho H, Akbarpour M, Delbandi AA. SLAM-family receptors come of age as a potential molecular target in cancer immunotherapy. Front Immunol. 2023;14:1174138. doi:10.3389/fimmu.2023.1174138

12. Dragovich MA, Mor A. The SLAM family receptors: Potential therapeutic targets for inflammatory and autoimmune diseases. Autoimmun Rev. Jul 2018;17(7):674–682. doi:10.1016/j.autrev.2018.01.018

13. Radhakrishnan SV, Bhardwaj N, Luetkens T, Atanackovic D. Novel anti-myeloma immunotherapies targeting the SLAM family of receptors. Oncoimmunology. 2017;6(5):e1308618. doi:10.1080/2162402X.2017.1308618

14. Yurchenko M, Skjesol A, Ryan L, et al. SLAMF1 is required for TLR4-mediated TRAM-TRIF-dependent signaling in human macrophages. J Cell Biol. Apr 2 2018;217(4):1411–1429. doi:10.1083/jcb.201707027

15. Nilsen KE, Zhang B, Skjesol A, et al. Peptide derived from SLAMF1 prevents TLR4-mediated inflammation in vitro and in vivo. Life Science Alliance. 2023;6(12):e202302164. doi:10.26508/lsa.202302164

16. Olsen MB, Kong XY, Louwe MC, et al. SLAMF1-derived peptide exhibits cardio protection after permanent left anterior descending artery ligation in mice. Brief Research Report. Frontiers in Immunology. 2024-April-15 2024;15doi:10.3389/fimmu.2024.1383505

17. Vrábel D, Pour L, Ševčíková S. The impact of NF-κB signaling on pathogenesis and current treatment strategies in multiple myeloma. Blood Reviews. 2019/03/01/ 2019;34:56–66. 10.1016/j.blre.2018.11.003

18. Akesolo O, Buey B, Beltrán-Visiedo M, Giraldos D, Marzo I, Latorre E. Toll-like receptors: New targets for multiple myeloma treatment? Biochem Pharmacol. May 2022;199:114992. doi:10.1016/j.bcp.2022.114992

19. Husebye H, Aune MH, Stenvik J, et al. The Rab11a GTPase controls Toll-like receptor 4-induced activation of interferon regulatory factor-3 on phagosomes. Immunity. Oct 29 2010;33(4):583–96. doi:10.1016/j.immuni.2010.09.010

20. Chesi M, Matthews GM, Garbitt VM, et al. Drug response in a genetically engineered mouse model of multiple myeloma is predictive of clinical efficacy. Blood. 2012;120(2):376–385. doi:10.1182/blood-2012-02-412783

21. Baranowska K, Misund K, Starheim KK, et al. Hydroxychloroquine potentiates carfilzomib toxicity towards myeloma cells. Oncotarget. Oct 25 2016;7(43):70845–70856. doi:10.18632/oncotarget.12226

22. Holien T, Vatsveen TK, Hella H, et al. Bone morphogenetic proteins induce apoptosis in multiple myeloma cells by Smad-dependent repression of MYC. Leukemia. May 2012;26(5):1073–80. doi:10.1038/leu.2011.263

23. Soriano GP, Besse L, Li N, et al. Proteasome inhibitor-adapted myeloma cells are largely independent from proteasome activity and show complex proteomic changes, in particular in redox and energy metabolism. Leukemia. Nov 2016;30(11):2198–2207. doi:10.1038/leu.2016.102

24. Nedal TMV, Moen SH, Roseth IA, et al. Diet-induced obesity reduces bone marrow T and B cells and promotes tumor progression in a transplantable Vk*MYC model of multiple myeloma. Sci Rep. Feb 13 2024;14(1):3643. doi:10.1038/s41598-024-54193-8

25. Wang Y, Muylaert C, Wyns A, et al. S-adenosylmethionine biosynthesis is a targetable metabolic vulnerability in multiple myeloma. Haematologica. 01/01 2024;109(1):256–271. doi:10.3324/haematol.2023.282866

26. Sarin V, Yu K, Ferguson ID, et al. Evaluating the efficacy of multiple myeloma cell lines as models for patient tumors via transcriptomic correlation analysis. Leukemia. 2020/10/01 2020;34(10):2754–2765. doi:10.1038/s41375-020-0785-1

27. Drexler HG, Matsuo Y. Malignant hematopoietic cell lines: in vitro models for the study of multiple myeloma and plasma cell leukemia. Leuk Res. Aug 2000;24(8):681–703. doi:10.1016/s0145-2126(99)00195-2

28. Klein B, Zhang XG, Lu ZY, Bataille R. Interleukin-6 in human multiple myeloma. Blood. Feb 15 1995;85(4):863–72.

29. Frassanito MA, Cusmai A, Iodice G, Dammacco F. Autocrine interleukin-6 production and highly malignant multiple myeloma: relation with resistance to drug-induced apoptosis. Blood. Jan 15 2001;97(2):483–9. doi:10.1182/blood.v97.2.483

30. Kawano Y, Fujiwara S, Wada N, et al. Multiple myeloma cells expressing low levels of CD138 have an immature phenotype and reduced sensitivity to lenalidomide. Int J Oncol. Sep 2012;41(3):876–84. doi:10.3892/ijo.2012.1545

31. Isoda A, Saito R, Komatsu F, et al. Palonosetron, aprepitant, and dexamethasone for prevention of nausea and vomiting after high-dose melphalan in autologous transplantation for multiple myeloma: A phase II study. Int J Hematol. Apr 2017;105(4):478–484. doi:10.1007/s12185-016-2152-6

32. Oancea M, Mani A, Hussein MA, Almasan A. Apoptosis of multiple myeloma. Int J Hematol. Oct 2004;80(3):224–31. doi:10.1532/ijh97.04107

33. Taabazuing CY, Okondo MC, Bachovchin DA. Pyroptosis and Apoptosis Pathways Engage in Bidirectional Crosstalk in Monocytes and Macrophages. Cell Chem Biol. Apr 20 2017;24(4):507–514 e4. doi:10.1016/j.chembiol.2017.03.009

34. Hideshima T, Anderson KC. Signaling Pathway Mediating Myeloma Cell Growth and Survival. Cancers (Basel). Jan 8 2021;13(2)doi:10.3390/cancers13020216

35. Tai Y-T, Li X-F, Breitkreutz I, et al. Inhibition of ERK1/2 Activity by the MEK1/2 Inhibitor AZD6244 (ARRY-142886) Induces Human Multiple Myeloma Cell Apoptosis in the Bone Marrow Microenvironment: A New Therapeutic Strategy for MM. Blood. 2006/11/16/ 2006;108(11):3460. 10.1182/blood.V108.11.3460.3460

36. Shaffer AL, Emre NCT, Lamy L, et al. IRF4 addiction in multiple myeloma. Nature. 2008/07/01 2008;454(7201):226–231. doi:10.1038/nature07064

37. Wang L, Yao ZQ, Moorman JP, Xu Y, Ning S. Gene expression profiling identifies IRF4-associated molecular signatures in hematological malignancies. PLoS One. 2014;9(9):e106788. doi:10.1371/journal.pone.0106788

38. Agnarelli A, Chevassut T, Mancini EJ. IRF4 in multiple myeloma-Biology, disease and therapeutic target. Leuk Res. Sep 2018;72:52–58. doi:10.1016/j.leukres.2018.07.025

39. Ohguchi H, Hideshima T, Bhasin MK, et al. The KDM3A-KLF2-IRF4 axis maintains myeloma cell survival. Nat Commun. Jan 5 2016;7:10258. doi:10.1038/ncomms10258

40. Chesi M, Robbiani DF, Sebag M, et al. AID-dependent activation of a MYC transgene induces multiple myeloma in a conditional mouse model of post-germinal center malignancies. Cancer Cell. Feb 2008;13(2):167–80. doi:10.1016/j.ccr.2008.01.007

41. Manning LS, Berger JD, O’Donoghue HL, Sheridan GN, Claringbold PG, Turner JH. A model of multiple myeloma: culture of 5T33 murine myeloma cells and evaluation of tumorigenicity in the C57BL/KaLwRij mouse. Br J Cancer. Dec 1992;66(6):1088–93. doi:10.1038/bjc.1992.415

42. Xie J, Bi Y, Zhang H, et al. Cell-Penetrating Peptides in Diagnosis and Treatment of Human Diseases: From Preclinical Research to Clinical Application. Front Pharmacol. 2020;11:697. doi:10.3389/fphar.2020.00697

43. Marqus S, Pirogova E, Piva TJ. Evaluation of the use of therapeutic peptides for cancer treatment. Journal of Biomedical Science. 2017/03/21 2017;24(1):21. doi:10.1186/s12929-017-0328-x

44. Lemech CR, Kichenadasse G, Marschner JP, Alevizopoulos K, Otterlei M, Millward M. ATX-101, a cell-penetrating protein targeting PCNA, can be safely administered as intravenous infusion in patients and shows clinical activity in a Phase 1 study. Oncogene. Feb 2023;42(7):541–544. doi:10.1038/s41388-022-02582-6

45. Shore N, Tutrone R, Roehrborn CG. Efficacy and safety of fexapotide triflutate in outpatient medical treatment of male lower urinary tract symptoms associated with benign prostatic hyperplasia. Ther Adv Urol. Jan-Dec 2019;11:1756287218820807. doi:10.1177/1756287218820807

46. Duan P, Ai L. Effect of TAK-242 on Proliferation and Apoptosis of Human Multiple Myeloma Cells. Cancer Research on Prevention and Treatment. 2019;46(4):322–326. doi:10.3971/j.issn.1000-8578.2019.18.1743

47. Nieto MJ, Hedjar A, Locke M, Caro J, Saif MW. Analysis of Updates in Multiple Myeloma Treatment and Management. J Clin Haematol. 2023;4(1):35–42. doi:10.33696/haematology.4.055

48. Wang C, Wang W, Wang M, et al. Different evasion strategies in multiple myeloma. Front Immunol. 2024;15:1346211. doi:10.3389/fimmu.2024.1346211

49. Painuly U, Kumar S. Efficacy of bortezomib as first-line treatment for patients with multiple myeloma. Clin Med Insights Oncol. 2013;7:53–73. doi:10.4137/cmo.S7764

50. Wallington-Beddoe CT, Sobieraj-Teague M, Kuss BJ, Pitson SM. Resistance to proteasome inhibitors and other targeted therapies in myeloma. Br J Haematol. Jul 2018;182(1):11–28. doi:10.1111/bjh.15210

51. Bayraktar UD, Bashir Q, Qazilbash M, Champlin RE, Ciurea SO. Fifty Years of Melphalan Use in Hematopoietic Stem Cell Transplantation. Biology of Blood and Marrow Transplantation. 2013/03/01/ 2013;19(3):344–356. 10.1016/j.bbmt.2012.08.011

52. McElwain TJ, Powles RL. High-dose intravenous melphalan for plasma-cell leukaemia and myeloma. Lancet. Oct 8 1983;2(8354):822–4. doi:10.1016/s0140-6736(83)90739-0

53. Attal M, Lauwers-Cances V, Hulin C, et al. Lenalidomide, Bortezomib, and Dexamethasone with Transplantation for Myeloma. N Engl J Med. Apr 6 2017;376(14):1311–1320. doi:10.1056/NEJMoa1611750

54. Gourzones-Dmitriev C, Kassambara A, Sahota S, et al. DNA repair pathways in human multiple myeloma: role in oncogenesis and potential targets for treatment. Cell Cycle. Sep 1 2013;12(17):2760–73. doi:10.4161/cc.25951

55. Agius MP, Hevenor L, Payne NC, et al. Pharmacological Targeting of IRF4 As a Therapeutic Strategy for Multiple Myeloma. Blood. 2024/11/05/ 2024;144:155. 10.1182/blood-2024-203381

56. Van Valckenborgh E, Matsui W, Agarwal P, et al. Tumor-initiating capacity of CD138- and CD138+ tumor cells in the 5T33 multiple myeloma model. Leukemia. Jun 2012;26(6):1436–9. doi:10.1038/leu.2011.373

57. Deng C, Lipstein MR, Scotto L, et al. Silencing c-Myc translation as a therapeutic strategy through targeting PI3Kdelta and CK1epsilon in hematological malignancies. Blood. Jan 5 2017;129(1):88–99. doi:10.1182/blood-2016-08-731240

