## Supplemental data for "Novel SLAMF1-derived peptide induces apoptosis in multiple myeloma cells by targeting IRF4 transcription factor for degradation"

### Supplementary methods

#### *Animal studies*

All animal experiments were performed in compliance with the ARRIVE guidelines, procedures approved by ethical committee. The relevant guidelines and regulations of animal handling and procedures for toxicity evaluation and C57BL6/Vk12653 mice model were approved by the Norwegian Food Safety Authority (FOTS ID 29802). *Toxicity evaluation.* Female 4-6 weeks C57BL/6 mice were provided by Janvier (France). Mice were acclimatized for 2 weeks at the pathogen free unit of Comparative Medicine Core facility (CoMed, NTNU, Trondheim, Norway). Mice were housed 6 mice/cage, randomly allocated to the two study groups: vehicle (sterile water) or 17 mg/kg P7N4-Pen that were injected intraperitoneally (i.p.) three times per week for 2 weeks. Injection volumes did not exceed 200 μl. Temperature was measured with an infrared thermometer for small rodents (BIO-IRB153, BioSeb Lab instruments, France). The animal's health was evaluated once a day by the personnel of animal facility. The health condition was monitored by using score sheets, the animals were weighed at least once a week. At day 14, mice were euthanized, and organs (spleen, liver and kidneys) were collected for macroscopic evaluation. Mice were anesthetized with isoflurane (5% isoflurane, 60% NO, 35% oxygen), bleed by cardiac puncture followed by cervical dislocation.

*C57BL6/Vk12653 MM model.* Female mice (6–8 weeks old, n = 24) were injected intravenously (i.v.) with $3 \times 10^5$ cells in 200 μl of PBS from a freshly thawed aliquot of bone marrow (BM) cells, which were previously isolated from tumor-bearing mice, with the aliquot containing 17% tumor cells. The Vk12653 cells were initially generously provided by Marta Chesi, Mayo Clinic [20]. 4 weeks after injection, mice were bled weekly from the saphenous vein to monitor M-component levels. Blood was collected to Eppendorf tubes, left at RT to coagulate for at least 30 min, centrifugated at 2000 x g for 10 min at RT and serum transferred to fresh tubes. Samples were diluted 1:80 in dialysis buffer. Serum M-component and albumin were measured using the urine procedure for protein electrophoresis on CAPILLARYS (Sebia). M-component was presented as a ratio of gamma globulin to albumin (G/A). 5 weeks after injection of cancer cells (day 36), animals with G/A ratio in the range 0.36-0.49 were allocated into 2 groups (n = 6 per group): vehicle (sterile water) or P7N4-Pen. Vehicle or 17 mg/kg P7N4-Pen were injected i.p. 4X/week for 2 weeks or until humane endpoints were reached. Injection volumes did not exceed 200 μl. On day 50 (7 weeks), mice were sacrificed. To evaluate the tumor load, mice spleens were harvested and weighed, and serum M-component (G/A ratio) was measured. Spleen cells were isolated by crushing the spleen in a cell culture dish containing 10 ml 5% FCS, RPMI1640 media, followed by passing the suspension through a 70 μm cell strainer. Cells were treated with RBC Lysis Buffer (Invitrogen), stained by live/dead stain (Invitrogen; Cat: L34957) according to the manufacturers protocol, blocked by of anti-mouse CD16/CD32 (Fc block) (Biolegend; Cat: 101302) (1 μg/million cells) on ice for 20 min, and stained on ice for 30 min with anti-mouse CD138 Brilliant Violet 421 (Biolegend; Cat: 142508), anti-mouse/human B220 Alexa Fluor® 647 (Biolegend; Cat: 103226) (1 μg/million cells) and anti-mouse CD19 PE/Cyanine7 (Biolegend; Cat: 115520) (2 μg/million cells) diluted in 100 μl of 0.1% BSA in PBS. Cells were analyzed by flow cytometry (LSRII flow cytometer, BD Biosciences, USA) to calculate the fraction of MM cells in the spleen.

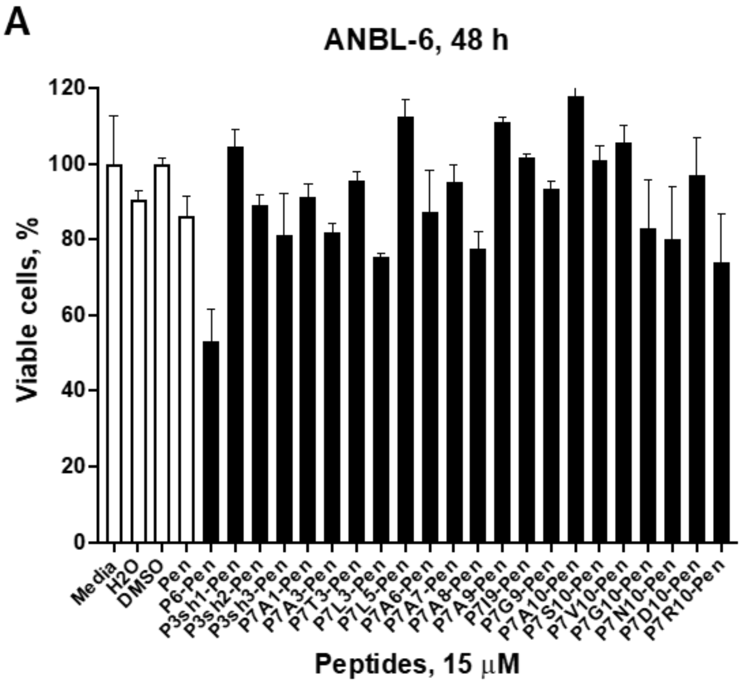


**B**

| Peptide | Sequence* |
|---|---|
| Pen | RQIKIWFQNRRMKWKK |
| P6-Pen | ITVYASVTLP G RQIKIWFQNRRMKWKK |
| P3sh1-Pen | TNSITVYASVTLT G RQIKIWFQNRRMKWKK |
| P3sh2-Pen | NSITVYASVTLT G RQIKIWFQNRRMKWKK |
| P3sh3-Pen | SITVYASVTLTG R QIKIWFQNRRMKWKK |
| P7A1-Pen | ATVYASVTLT G RQIKIWFQNRRMKWKK |
| P7A3-Pen | ITAYASVTLT G RQIKIWFQNRRMKWKK |
| P7T3-Pen | ITTYASVTLT G RQIKIWFQNRRMKWKK |
| P7L3-Pen | ITLYASVTLT G RQIKIWFQNRRMKWKK |
| P7L5-Pen | ITVYLSVTLT G RQIKIWFQNRRMKWKK |
| P7A6-Pen | ITVYAAVTLT G RQIKIWFQNRRMKWKK |
| P7A7-Pen | ITVYASATLT G RQIKIWFQNRRMKWKK |
| P7A8-Pen | ITVYASVALT G RQIKIWFQNRRMKWKK |
| P7A9-Pen | ITVYASVTAT G RQIKIWFQNRRMKWKK |
| P7I9-Pen | ITVYASVTIT G RQIKIWFQNRRMKWKK |
| P7G9-Pen | ITVYASVTGT G RQIKIWFQNRRMKWKK |
| P7A10-Pen | ITVYASVTAA G RQIKIWFQNRRMKWKK |
| P7S10-Pen | ITVYASVTLS G RQIKIWFQNRRMKWKK |
| P7V10-Pen | ITVYASVTLV G RQIKIWFQNRRMKWKK |
| P7G10-Pen | ITVYASVTLG G RQIKIWFQNRRMKWKK |
| P7N10-Pen | ITVYASVTLN G RQIKIWFQNRRMKWKK |
| P7D10-Pen | ITVYASVTLD G RQIKIWFQNRRMKWKK |
| P7R10-Pen | ITVYASVTLR G RQIKIWFQNRRMKWKK |

*Glycine amino acid is a linker between peptide and Pen

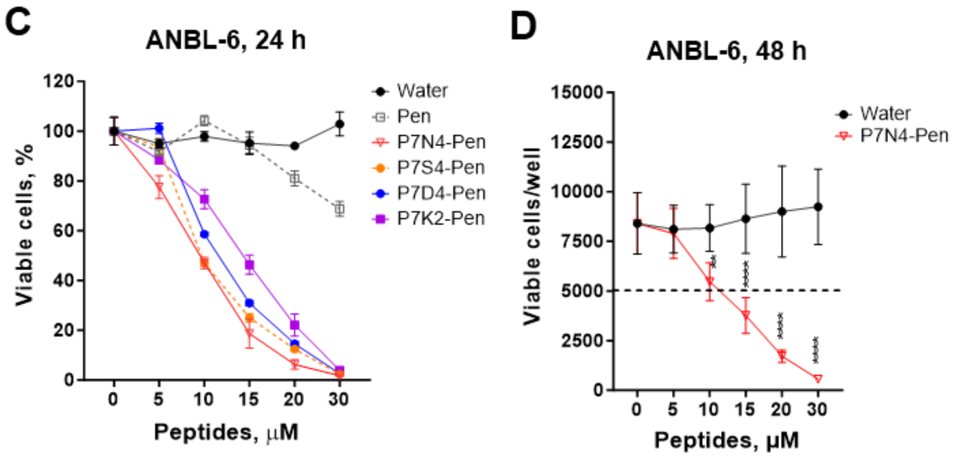


**Supplementary Fig. 1. P7-Pen variants with single amino acid substitutions other than at positions 2 and 4 have no effect on ANBL-6 cell viability.** (**A**) ANBL-6 cells were grown in normal media, or treated with solvents (water, DMSO), 15 μM control peptide Pen or P7 peptide variants (**B**) for 48 h before measuring cell viability (3 independent experiments). (**B**) Table with amino acid sequences for SLAMF1-derived peptides. Amino acid substitutions in relation to P7-Pen peptide are highlighted in red. (**C**) ANBL-6 cells were treated with increasing concentrations (range of 5-30 μM) of peptide or control treatments (water, Pen) for 24 h prior cell viability analysis. (**D**) The graph presents the values of viable cells per well, calculated for water and P7N4-Pen treatments from Figure 1C using the standard curve for cells per well. The dashed line represents the number of cells per well at the time of seeding. (**A, C, D**) Cell viability was measured using the CellTiterGlo assay. Data presented as the mean percentage of viable cells to the media control (100%), with mean ± SD for three technical replicates (**C**) or mean ± SEM for three biological replicates (**A, D**). Significance evaluated using (**A**) one-way ANOVA, (**D**) two-way ANOVA, significance levels: *P**<0.05, *P***<0.01, *P****<0.001, *P*****<0.0001).

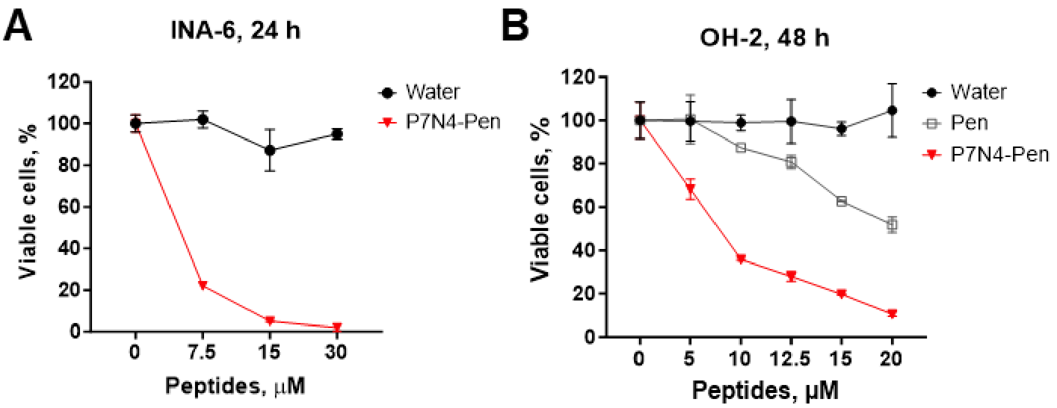


**Supplementary Fig. 2. Pilot screening of P7N4-Pen's effect on the viability of additional MM cell lines.** IL-6 dependent INA-6 (**A**) and OH-2 cells (**B**) have got P7N4-Pen or control treatments (water or control peptide Pen) and incubated for 48 h before measuring cell viability with the CellTiterGlo assay. Data presented as the mean ± SD percentage of viable cells when related to the water control (100%) for three biological replicates.

A

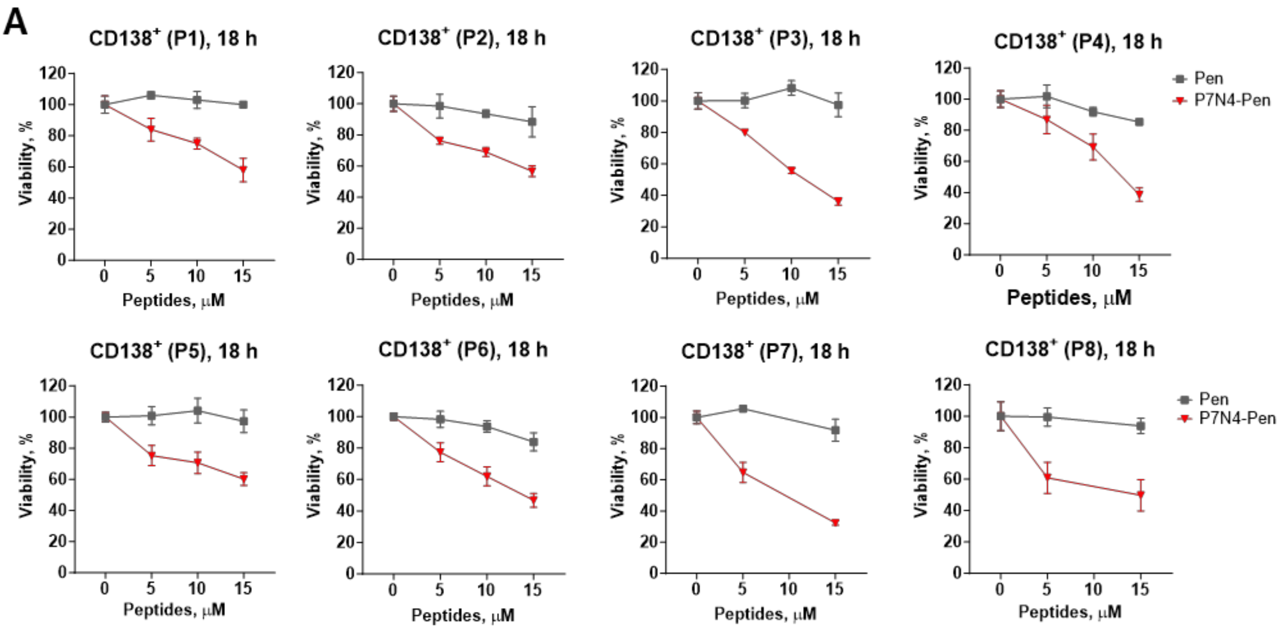


| Patient | P1 | P2 | P3 | P4 | P5 | P6 | P7 | P8 |
|---|---|---|---|---|---|---|---|---|
| $IC_{40}$ (µM) | 14.6 | 13.8 | 9.1 | 12 | 15 | 11.2 | 7.2 | 7.2 |

B

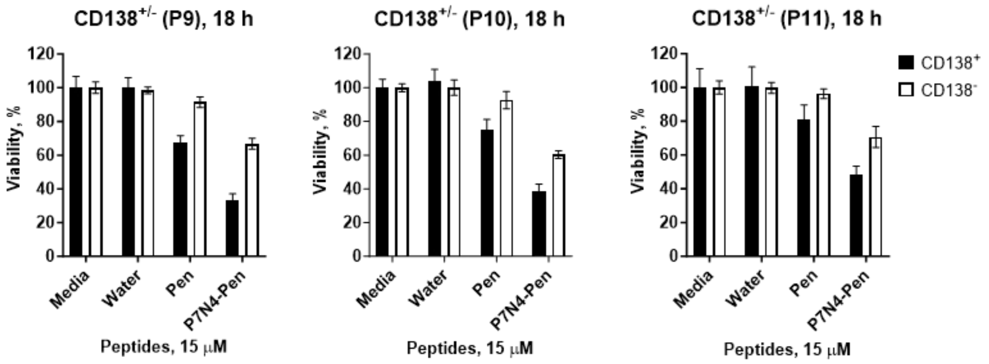


**Supplementary Fig. 3. P7N4-Pen peptide induces cell death in primary MM CD138$^{+}$ cells with patient-dependent variability and has a lesser impact on CD138$^{-}$ cell viability.** (**A**) Graphs depict the dose-dependent viability of CD138$^{+}$ primary MM cells from individual patients (n = 8) treated with either the control peptide Pen or P7N4-Pen for 18 h. $IC_{40}$ values (µM) for each patient are shown in the table below. (**B**) Cells were incubated in media, media with water, or 15 µM peptides for 18 h before cell viability was assessed. Graphs depict the viability of primary bone marrow CD138$^{+}$ and CD138$^{-}$ cells, with a separate graph for each patient (n = 3). (**A, B**) Cell viability was assessed using the CellTiter-Glo assay. Data are presented as mean ± SD (%) of viable cells relative to the media control (100%) from three technical replicates.

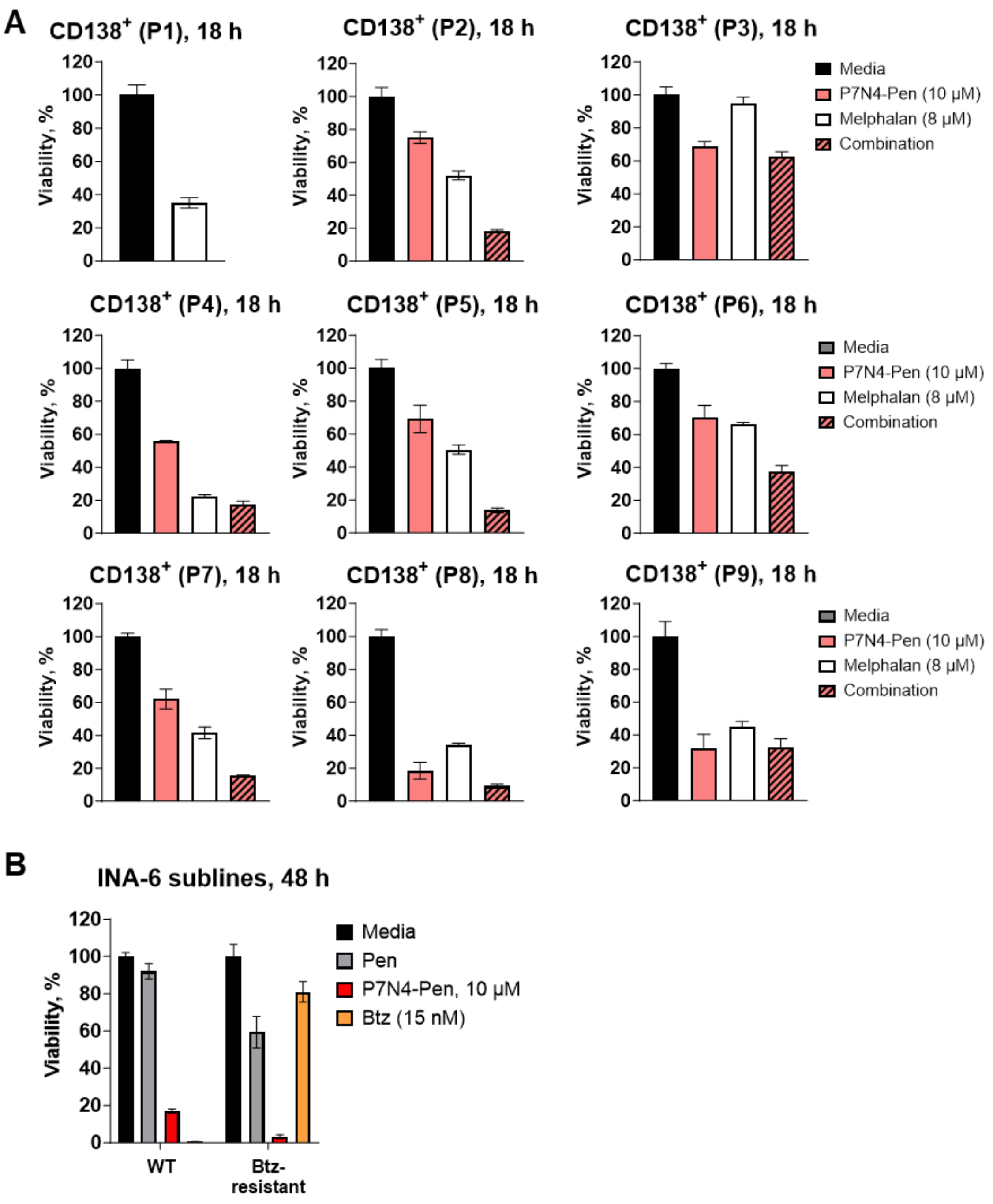


**Supplemental Fig. 4. P7N4-Pen enhances melphalan-induced reduction in CD138$^+$ primary MM cell viability and decreases the viability of IL-6-dependent INA-6 cells resistant to bortezomib.** (**A**) Viability analysis of primary CD138$^+$ MM cells for individual MM patients (n = 9), which supplements combined graph presented on Fig. 4C. Cells were treated for 18 h with 8 μM melphalan or 10 μM P7N4-Pen, or combination. Data presented as mean ± SD for technical replicates. (**B**) Wild type (WT) INA-6 cells or INA-6 subline resistant to bortezomib (Btz) were treated for 48 h with 10 μM peptides or 15 nM Btz, followed by measuring cell viability. Data presented as mean ± SD for two biological replicates. In all assays cell viability was evaluated using the CellTiter-Glo assay.

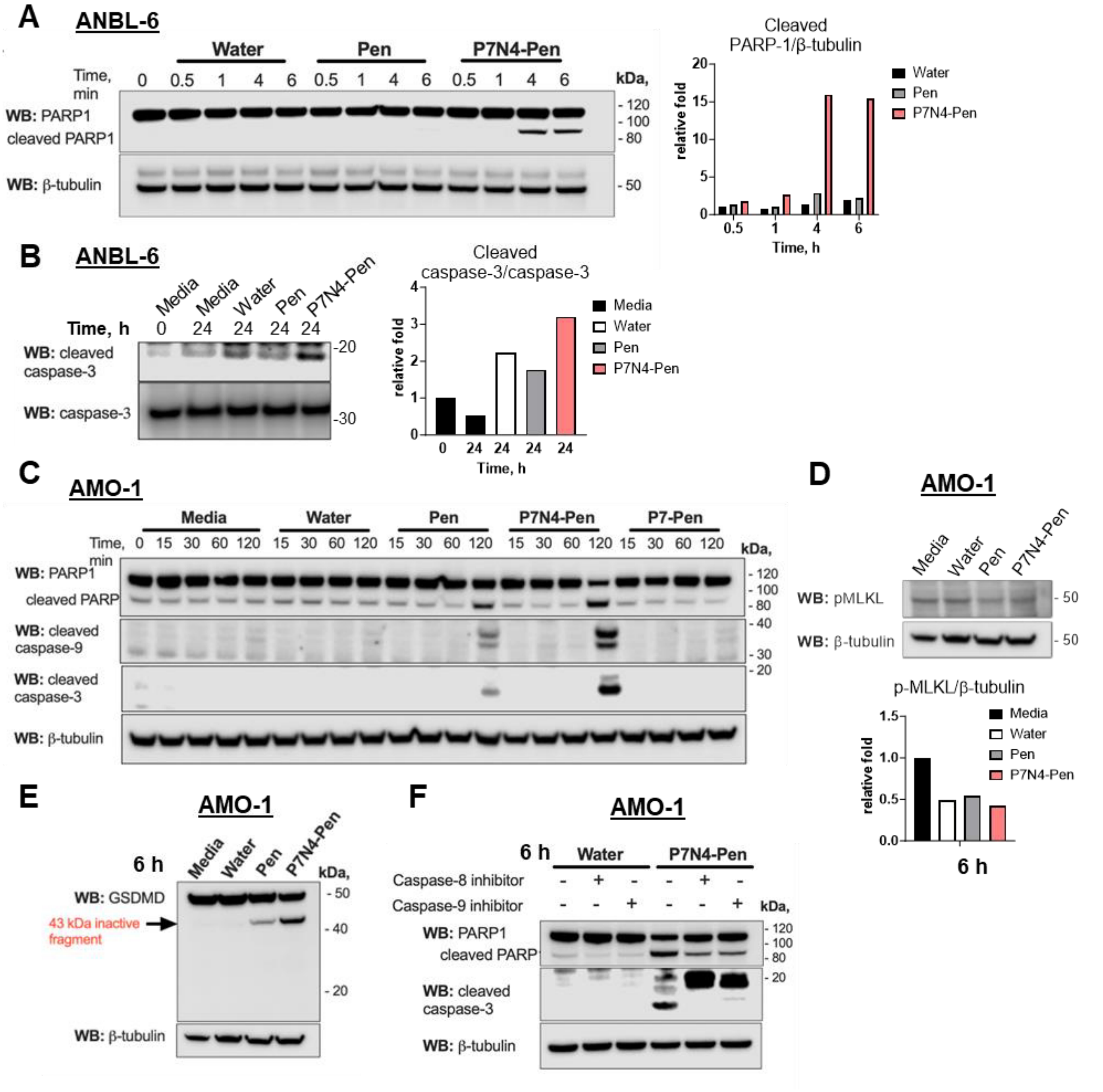


**Supplementary Fig. 5. P7N4-Pen induces cell death through caspase-8- and caspase-9-dependent apoptosis, with no contribution from pyroptosis or necroptosis.** (**A, B**) Representative image (n = 3) for WB analysis of PARP cleavage (**A**) or caspase-3 cleavage (**B**) in ANBL-6 lysates pre-treated with peptides (10 μM) or control treatment (water) for the indicated time. Graphs show the relative fold for cleaved PARP-1 to β-tubulin (**A**) or cleaved to uncleaved caspase-3 (**B**). (**C**) Full size image with all treatments tested for the main Figure 4D. (**D, E**) AMO-1 cells were pre-treated for 6 h by 7.5 μM peptides or control treatment, followed by WB analysis of MLKL Ser358 phosphorylation (**D**) or GSDM cleavage (**E**). (**D**) The relative fold for pMLKL level to β-tubulin for the representative image is shown on graph. (**F**) AMO-1 cells were pre-treated with caspase-8 or caspase-9 inhibitor (40 nM) prior addition of 7.5 μM peptides or control treatments. (**A, C-F**) WB for total β-tubulin was applied for loading control. (**D-E**) Representative images from two independent experiments are shown.

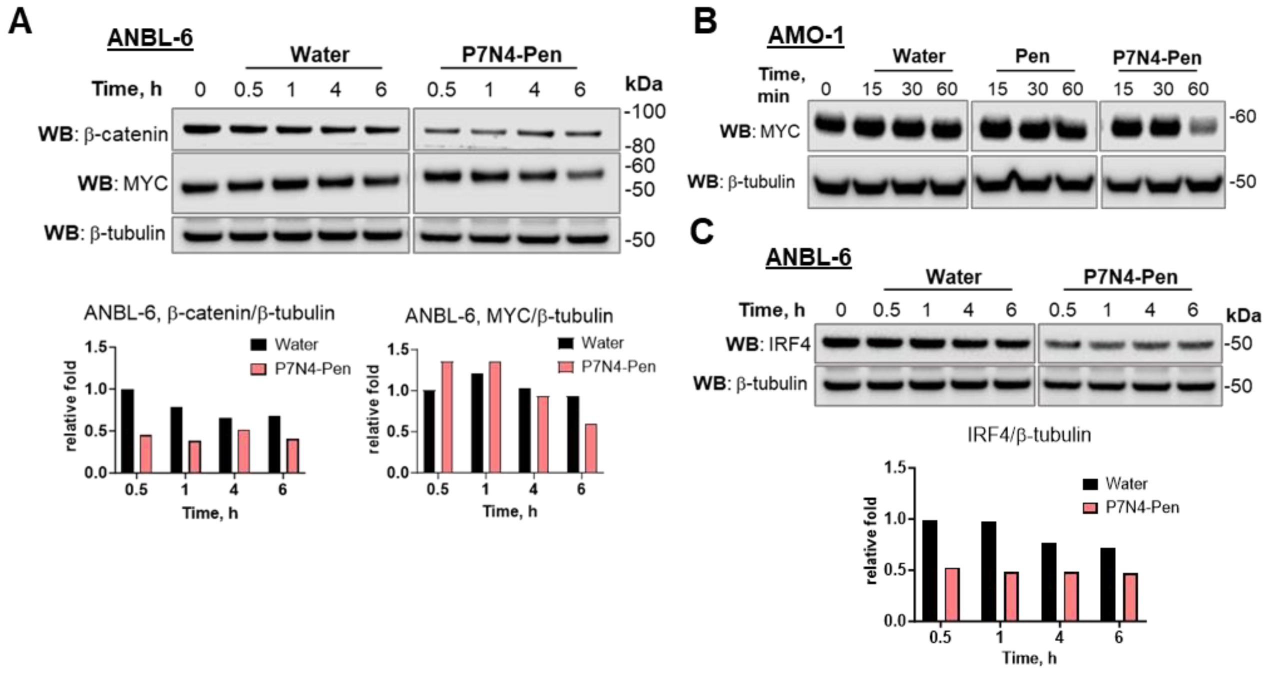


**Supplementary Fig. 6. P7N4-Pen downregulates IRF4, MYC, and β-catenin protein expression in AMO-1 and ANBL-6 HMCLs.** (**A**) ANBL-6 cells were treated with 10 µM peptides, followed by WB analysis for total β-catenin and MYC. Graphs show the relative fold of β-catenin or MYC level to β-tubulin calculated for representative images from three independent experiments. (**B, C**) AMO-1 and ANBL6 cells were treated with 7.5 µM or 10 µM peptides, respectively, followed by WB analysis of IRF4 or MYC protein expression in cell lysates. (**C**) Graph depicts the relative fold of IRF4 to β- calculated for representative images from three independent experiments. (**A-C**) Images shown for Western blots for the selected conditions/treatments are taken from the same membranes.

.

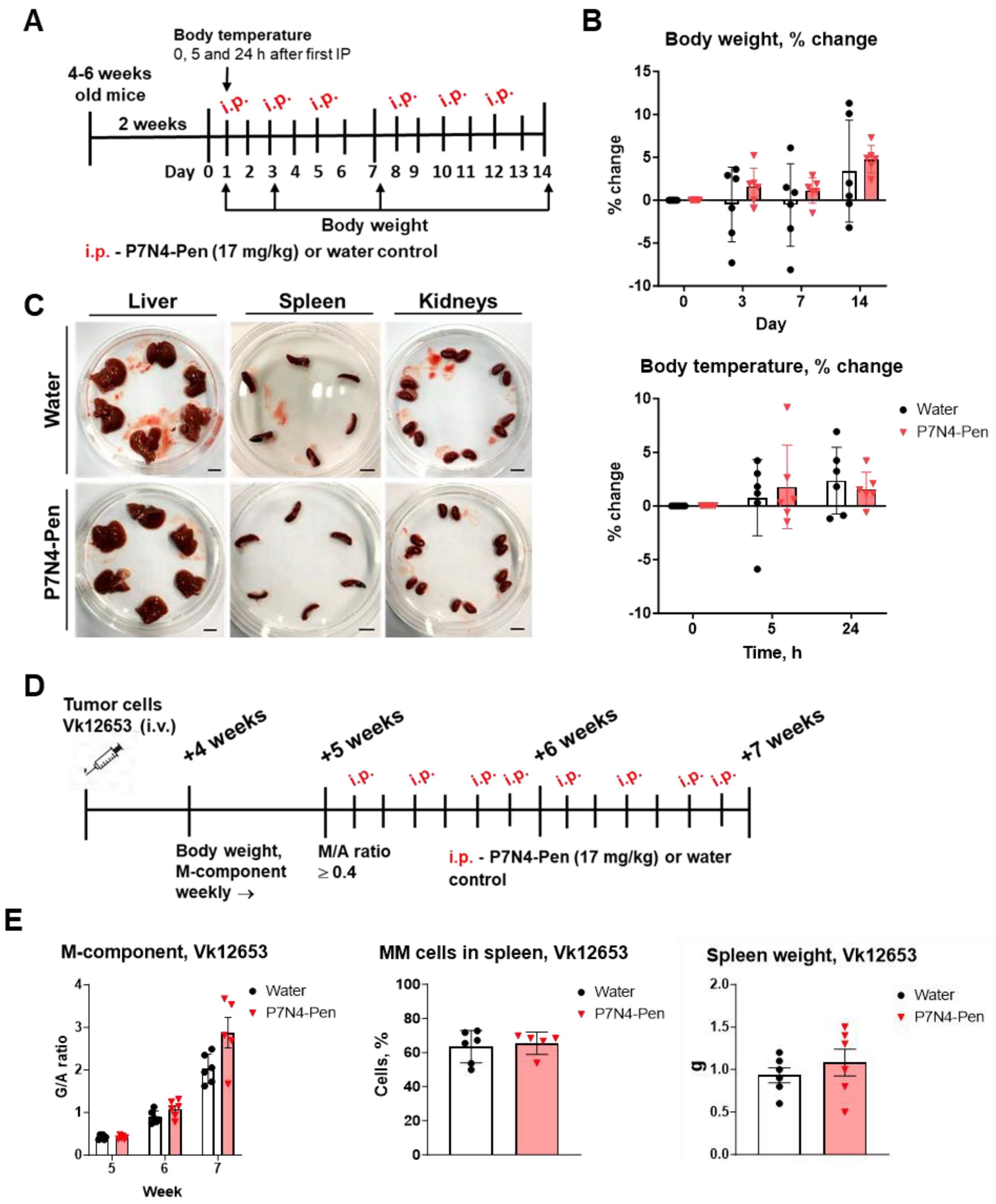


**Supplementary Fig. 7. P7N4-Pen is well tolerated by mice but does not affect Vk12653 tumor growth *in vivo.*** (**A**) Graphical overview of the setup and timeline for the peptide toxicity evaluation in healthy C57BL/6 mice. Mice (4–6 weeks old) were acclimatized in the animal facility for 14 days before injections began. Mice received intraperitoneal (i.p.) injections of either peptide solvent (water) or 17 mg/kg P7N4-Pen, administered seven times over a 2-week period before euthanasia. (**B**) Graphs are showing mice body weight (top) on days 0, 3, 7 and 14 and body temperature (bottom) before first injection, and at 5 h and 24 h after the first injection. Data presented as mean with SEM (n = 6 per group). (**C**) Images of livers, spleens and kidneys of all C57BL/6 mice from the toxicity test experiment that were used for macroscopic evaluation, scale bar 1 cm. (**D**) Graphical overview of the setup and timeline for Vk12653 murine MM. C57BL/6 mice (n = 36) were injected intravenously (i.v.)

with Vk12653 cells. After 4 weeks, mice were bled from the saphenous vein to monitor the serum M-component. 5 weeks after injection of cancer cells, mice with G/A ratio in the range 0.35-0.5 (n = 18) were allocated to three treatment groups (6 mice/group) prior to starting i.p. injection of water (control) or 17 mg/kg P7N4-Pen four times per week. Mice were euthanized 7 weeks after injection of cancer cells or when humane endpoints were reached. (**E**) Serum M-component is presented as globulin to albumin (G/A) ratio) (left panel). Percentage of MM tumor cells at the endpoint in spleens of Vk12653 mice was established by staining of isolated spleen cells with anti-CD138, anti-B220 and anti-CD19 Abs and flow cytometry (middle panel). Spleen weight was measured before isolation of cells at endpoint (right panel). Data presented as mean ± SEM. Significance evaluated using (**B**) two-way ANOVA, Dunnett's multiple comparisons test; (**E**) two-way ANOVA, Sidaks multiple comparisons test; i.v.- intravenous, i.p.- intraperitoneal.

**Supplementary Table 1**. Differential expression of IRF4-associated signature genes and *IRF4, MYC* and *KLF2**

| | AMO-1 | | RPMI-8226 | |
|---|---|---|---|---|
| | P7N4-Pen to media control | | P7N4-Pen to media control | |
| **gene** | **log2fold** | **Padj** | **log2fold** | **Padj** |
| *ACVR1B* | -1.203792 | 2.46E-16 | -0.25626079 | 0.1036783 |
| *CASP10* | -1.30832 | 4.798E-27 | -1.14215219 | 7.587E-20 |
| *CDKN1A* | 0.9859125 | 1.097E-05 | 1.715732219 | 9.201E-14 |
| *CITED2* | 1.0521858 | 2.306E-11 | -1.24077158 | 8.363E-15 |
| *DUSP1* | 3.1023328 | 2.871E-12 | 0.065994253 | 0.8277767 |
| *DUSP5* | 3.997016 | 2.28E-39 | -0.25452085 | 0.2745965 |
| *ESR1* | 0.7105665 | 2.347E-05 | 0.152073682 | 0.5660033 |
| *FOSB* | 7.31383 | 1.296E-40 | 4.322181907 | 1.425E-14 |
| *JUN* | 3.6825118 | 2.305E-15 | 0.262728756 | 0.2973848 |
| *JUNB* | 2.018885 | 5.625E-09 | 0.284516826 | 0.2424059 |
| *KANK1* | -1.346284 | 1.514E-26 | -0.02610718 | 0.8969769 |
| *SDC1* | -0.644626 | 1.596E-09 | 0.152263981 | 0.2094366 |
| *SRGN* | 1.962575 | 1.475E-76 | -0.02479909 | 0.8878465 |
| *TXNIP* | -3.295408 | 1.347E-42 | -1.84775066 | 1.814E-14 |
| *IRF4* | -0.130486 | 0.685653 | -0.11906388 | 0.6292994 |
| *KLF2* | -2.772585 | 9.632E-15 | -0.44223979 | 0.0789094 |
| *MYC* | -2.636032 | 7.877E-24 | -0.00135432 | 1 |

*AMO-1 and RPMI-8226 cells were treated with 10 μM P7N4-Pen or media for 90 min prior to RNA isolation and RNA sequencing, with three biological replicates for each condition. Table contains data for $log_2$fold and Padj for the differential analysis of expression of *IRF4, KLF2, MYC* and IRF4-associated molecular signature genes [37], used for generation of heatmap in Fig. 6G. Genes for IRF4-associated signature were selected based on the differential analysis of their expression in AMO-1 cells by pairwise analysis for P7N4-Pen *vs.* media samples, $log_2$fold change > 0.4, $log_2$fold change < -0.4, Padj < 0.05).

**Supplementary Table 2**. Z-transformed $log_2$TPM values for differentially expressed IRF4-associated signature genes, *IRF4, MYC* and *KLF2**

| | **AMO-1** | | | | | | **RPMI-8226** | | | | | |
|---|---|---|---|---|---|---|---|---|---|---|---|---|
| | **media** | | | **P7N4-Pen peptide** | | | **media** | | | **P7N4-Pen peptide** | | |
| **Gene** | **1** | **2** | **3** | **1** | **2** | **3** | **1** | **2** | **3** | **1** | **2** | **3** |
| *CASP10* | 1.135 | 0.985 | 0.916 | -0.846 | -0.830 | -0.894 | -0.024 | 0.476 | -0.112 | -1.443 | -2.041 | -1.184 |
| *SDC1* | 1.440 | 1.540 | 1.460 | 1.013 | 1.199 | 1.126 | -0.950 | -0.937 | -1.019 | -0.798 | -0.905 | -0.906 |
| *DUSP5* | 0.137 | -0.265 | -0.197 | 2.061 | 2.180 | 2.118 | -0.092 | -0.011 | -0.334 | -0.151 | -0.488 | -0.611 |
| *KANK1* | -0.104 | -0.234 | -0.083 | -1.109 | -1.022 | -1.005 | -0.816 | -0.839 | -0.765 | -0.672 | -0.936 | -0.908 |
| *CDKN1A* | 0.499 | 0.739 | 1.001 | 1.408 | 1.831 | 1.588 | -1.861 | -1.462 | -2.155 | -0.139 | -0.317 | -0.571 |
| *ACVR1B* | 1.593 | 1.743 | 1.883 | -0.372 | -0.847 | -0.124 | -0.976 | -0.863 | -0.703 | -1.756 | -1.802 | -0.795 |
| *DUSP1* | -0.594 | -1.226 | -0.641 | 1.923 | 1.973 | 1.963 | -0.066 | 0.375 | 0.485 | 0.357 | 0.867 | -0.069 |
| *JUN* | -1.956 | -2.128 | -1.810 | 0.985 | 1.335 | 0.646 | -0.409 | -0.435 | -0.129 | -0.217 | 0.614 | 0.120 |
| *FOSB* | -0.651 | -0.681 | -0.724 | 2.005 | 1.759 | 2.064 | -1.241 | -0.802 | -0.952 | 0.572 | 0.866 | 0.472 |
| *JUNB* | 0.708 | 0.389 | 0.468 | 2.329 | 2.192 | 2.306 | -1.205 | -0.894 | -0.998 | -0.527 | -0.482 | -0.673 |
| *SRGN* | -0.854 | -0.967 | -1.033 | 1.546 | 1.594 | 1.475 | -0.968 | -0.918 | -1.149 | -1.022 | -1.031 | -1.094 |
| *KLF2* | 1.408 | 1.111 | 0.948 | -1.057 | -0.592 | -1.161 | -0.409 | -0.183 | -0.793 | -0.949 | -0.735 | -1.534 |
| *MYC* | 1.005 | 0.907 | 0.947 | -0.418 | -0.490 | -0.396 | 0.851 | 0.846 | 0.810 | 0.838 | 0.966 | 0.671 |
| *TXNIP* | 0.759 | 1.246 | 0.754 | -1.766 | -1.764 | -2.089 | 1.230 | 1.423 | 1.320 | -0.216 | -0.859 | -0.050 |
| *CITED2* | -0.794 | -1.248 | -1.091 | 0.244 | 1.172 | 0.131 | 1.158 | 1.534 | 1.541 | -0.179 | -0.747 | -0.569 |
| *ESR1* | 0.467 | 0.388 | 0.470 | 0.807 | 0.668 | 0.691 | -0.953 | -2.203 | -1.522 | -1.015 | -1.417 | -1.221 |
| *IRF4* | 0.675 | 0.424 | 0.603 | 0.039 | 1.034 | -0.339 | -0.916 | -0.292 | -2.177 | -0.822 | -1.588 | -1.548 |

*Values used for preparation of heatmap shown on Figure 7A.

**Supplementary Table 3**. Normalized counts for *IRF4 and MYC* genes expression by AMO-1 and RPMI-8226 cells treated by P7N4-Pen or control treatments.

| Gene | Condition/ Replicate | AMO-1 | | | RPMI-8226 | | |
|---|---|---|---|---|---|---|---|
| | | **1** | **2** | **3** | **1** | **2** | **3** |
| ***IRF4*** | **Media** | 34199.2 | 31210.5 | 35480.53 | 21649.66 | 15732.74 | 8260.318 |
| | **Water** | 32663.04 | 26905.47 | 25998.78 | 19056.45 | 11810.48 | 12095.6 |
| | **P7N4-Pen** | 25630.32 | 42626.47 | 21133.7 | 11162.58 | 16508.38 | 11390.84 |
| ***MYC*** | **Media** | 32830.18 | 31086.89 | 35528.84 | 28635.36 | 28811.29 | 27248.95 |
| | **Water** | 20143.99 | 28443.91 | 18639.33 | 31369.38 | 40473.11 | 24520.67 |
| | **P7N4-Pen** | 5104.379 | 4631.816 | 5263.095 | 33698.78 | 28307.56 | 22538.34 |
